# Creatine synthesis is a tumor suppressor pathway hypostatic to one-carbon metabolism

**DOI:** 10.1101/2025.05.23.655758

**Authors:** Nicole Yong Ting Leung, Tiffany Melanie Yee, Roya AminiTabrizi, Ragini Bhalla, Nicholas Ang, Liangjian Lu, Chang Yien Chan, Derrick Wen Quan Lian, Matthew Y. Lim, Joao A. Paulo, Wei Wu, Hardik Shah, Liang Wei Wang

## Abstract

Methylene tetrahydrofolate reductase 2 (MTHFD2), the rate-limiting enzyme of mitochondrial one-carbon metabolism, is one of the most highly expressed metabolic enzymes across diverse cancers and lymphoproliferative disorders. However, its exact roles in oncogenic metabolism remain poorly defined. We show that MTHFD2 is a key regulator of mitochondrial energetics in Epstein-Barr virus-transformed B lymphoblastoid cell lines (LCLs), an *in vitro* model of post-transplant lymphoproliferative disorder (PTLD). We also delineate a role for MTHFD2 in fueling *de novo* creatine synthesis; MTHFD2 mediates the production of glycine, a necessary substrate for creatine synthesis, through serine catabolism. Aminomethyltransferase (AMT) suppression short-circuits the glycine cleavage system (GCS) to augment LCL mitochondrial glycine levels. Creatine synthesis is hypostatic to mitochondrial one-carbon metabolism; inhibition of creatine synthesis improves LCL fitness only when MTHFD2 is lost. Our findings emplace MTHFD2 at the nexus of amino acid and energy metabolism pathways in LCLs, with potential clinical ramifications for PTLD.

**Highlights:** • Complete activation of creatine synthesis in an *in vitro* cellular model of PTLD

• Creatine synthesis is a major sink for mitochondrial 1C-derived glycine

• Reverse GCS activity due to AMT deficiency in lymphoblastoid cells

• Epistasis between mitochondrial 1C metabolism and creatine synthesis

**eTOC Blurb:** Leung et al. demonstrate that MTHFD2 is crucial for creatine synthesis in lymphoproliferative disorders. MTHFD2 supports forward 1C flux through SHMT and drives reverse GCS activity to augment mitochondrial glycine, a substrate for creatine synthesis. Tumor-suppressive effects of creatine synthesis are unmasked with MTHFD2 loss, exhibiting metabolic epistasis.

## Introduction

Post-transplant lymphoproliferative disorder (PTLD) is a life-threatening complication following solid organ transplantation (SOT). In fact, PTLD is the most common form of cancer-associated lethality to arise after SOT in both pediatric and adult settings^1^. The condition arises when tissue-resident transformed lymphocytes are transferred during SOT into immunosuppressed transplant recipients, allowing the transformed cells to evade detection by the recipient’s immune system and develop into malignant growths. A key driver of PTLD is Epstein-Barr virus (EBV), the first described human cancer virus; the virus maintains latency in immunocompetent persons but can drive sustained growth and proliferation of the host cell e.g., donor lymphocytes that are inadvertently carried over in the transplanted organ or *de novo* infection of recipient’s lymphocytes, under conditions of immunosuppression, histo-incompatibility and recipient seronegativity towards EBV. Hence, to model PTLD disease-driving cells *in vitro*, EBV is used to transform resting B cells into continually dividing lymphoblastoid cell lines (LCLs) that have many molecular similarities to their *in vivo* counterparts. EBV-transformed LCLs activate pro-survival and growth-promoting signaling pathways and remodel cellular chromatin to install an oncogenic program. In previous work done by us^2^ and others^3^, LCL outgrowth is strongly influenced by mitochondrial folate-mediated one-carbon metabolism (FOCM).

Mitochondrial FOCM is a critical pathway that facilitates the generation of formate (the one-carbon unit) and glycine for nucleotide, protein and glutathione syntheses, as well as reducing equivalents. Under normal physiological conditions, FOCM is rarely upregulated, except in embryonic development^4^ and the expansion of activated lymphocytes^5,6^. Notably, apart from LCLs, this pathway is specifically and widely upregulated in cancers and lymphoproliferative diseases but not in healthy proliferating cells^7^, thus receiving renewed clinical and scientific interest in recent years as a specific target for therapeutic intervention.

Methylene tetrahydrofolate dehydrogenase 2 (MTHFD2) is a major enzyme in the mitochondrial branch of the FOCM pathway. It mediates the conversion of 5,10 methylenetetrahydrofolate (THF) to 10-formyl-THF, a key carrier of 1C units (Figure 1A). Apart from its enzymatic activity, MTHFD2 also has reported roles in regulating RNA splicing^8,9^, which may partly explain why its loss cannot be rescued by downstream metabolites, namely glycine and formate. In some cells, MTHFD2 loss can be well compensated by flux reversal through the cytosolic branch of the FOCM pathway^10^. Fundamentally, the mechanisms by which MTHFD2 promotes growth and proliferation of transformed cells have yet to be fully delineated.

**Figure 1.**
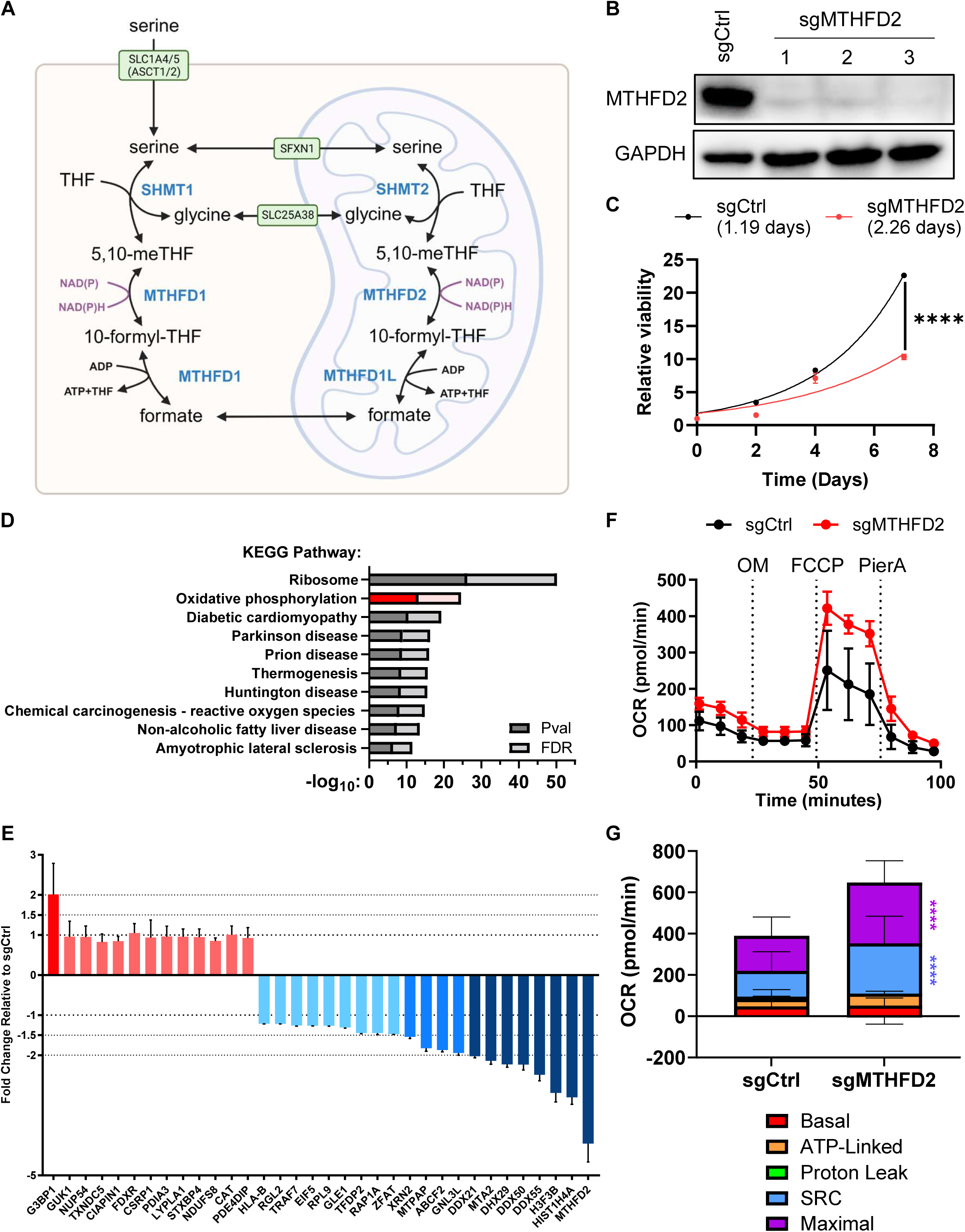
MTHFD2 regulates B lymphoblastoid cell line viability and energetics. (A) Schematic illustrating the one-carbon (1C) metabolic pathway. Enzymes, transporters, metabolites and redox-active pairs are indicated in blue, with green boxes, in black and in purple, respectively. (B) Representative immunoblot (N = 3) of cell lysates obtained from control (sgCtrl) and MTHFD2-targeting (sgMTHFD2) sgRNA-expressing GM12878 Cas9 cells. GAPDH was used as housekeeping control. (C) Growth curve analysis of GM12878 sgCtrl and sgMTHFD2 cells with calculated doubling times indicated. (D) KEGG analysis of transcriptomes from GM12878 sgCtrl and sgMTHFD2 cells. The top 10 enriched pathways are shown. Nominal P-values and false discovery rates (FDR) are plotted. The KEGG term “Oxidative phosphorylation” is highlighted in red and pink, instead of dark grey and light grey. (E) Proteomic analysis of GM12878 sgCtrl and sgMTHFD2 cells. Red: proteins significantly upregulated (P < 0.05) in sgMTHFD2 cells. Blue: proteins significantly downregulated (P < 0.05) in sgMTHFD2 cells. (F) Seahorse XF Cell Mito Stress Test on GM12878 sgCtrl and sgMTHFD2 cells. Oxygen consumption rates (OCR) were measured in response to mitochondrial poisons. OM, oligomycin; FCCP, carbonyl cyanide 4-(trifluoromethoxy)phenylhydrazone; PierA, piericidin A. (G) Respiratory parameters of GM12878 sgCtrl and sgMTHFD2 cells computed from OCR measurements from the Mito Stress Test. SRC, spare respiratory capacity. All quantitative data represent N = 3 biological replicates with errors bars indicative of SD. ***, p < 0.001; ****, p < 0.0001 by Student’s t-test (Panels C and G). See also Figure S1.

The mammalian phosphagen system, centered on creatine, has an important role in maintaining energy balance in tissues. Creatine and creatine phosphate (also known as phosphocreatine) act as important buffers for intracellular ATP in tissues facing elevated and fluctuating rates of energy demand, with *de novo* creatine synthesis accounting for approximately half the daily creatine requirement in a healthy adult. This pathway involves two enzymes – arginine-glycine amidinotransferase (AGAT; also known as GATM) and guanidinoacetate methyltransferase (GAMT). GATM mediates the condensation of glycine and arginine to produce guanidinoacetate (GAA) and ornithine, while GAMT mediates S-adenosylmethionine (SAM)-dependent methylation of GAA to produce creatine. In mice, some tissues e.g., liver and testis, co-express the creatine synthesis enzymes^11^, although it is not entirely clear whether the two enzymes are segregated into sub-anatomical niches and/or distinct specialized cells. Oligodendrocytes in the mouse brain can express both enzymes. In contrast, in humans, the pathway is thought to be separated into two distinct sites, namely the kidney and the liver, which express high levels of GATM and GAMT, respectively. Creatine is subsequently released into circulation to be taken up by tissues, especially skeletal muscle, for substrate-level phosphorylation to form phosphocreatine, a ready source of high-energy phosphates that can be rapidly mobilized for biochemical reactions in the cell.

In this study, we aim to address the gap in knowledge pertaining to MTHFD2’s role in LCL biology, particularly how it supports proliferation and survival. By using multi-omics approaches to characterize LCLs genetically edited to lack MTHFD2, we observed significant reprogramming of cells towards a hyper-oxidative phenotype, concomitant with drastic reductions in expansion potential and cell viability. Intracellular creatine was noticeably depleted in MTHFD2-deficient cells. Interestingly, either targeted disruption of GATM or supraphysiological supplementation of creatine (which acts as a pathway end-product inhibitor) restored MTHFD2-null population expansion and mitigated apoptosis, demonstrating the tumour suppressive character of the creatine synthesis pathway.

## Results

### MTHFD2 regulates B lymphoblastoid cell line viability and energetics

We first employed clustered regularly interspaced short palindromic repeats (CRISPR)/Cas9 technology to establish stable cell lines lacking MTHFD2 protein expression^12^ (Figure 1B) and confirmed that MTHFD2-deficient cells were expanding at a lower rate than their wild-type counterparts (Figure 1C). Non-linear curve fitting and subsequent analysis revealed that MTHFD2-deficient cells had a calculated doubling time of 2.26 days, which was approximately 90% longer than that of MTHFD2-sufficient cells (1.19 days). This agrees with previous reports which showed that MTHFD2 loss slowed down population expansion, rather than completely abolishing growth and proliferation^7^.

Messenger RNAs were isolated from control and MTHFD2-deficient cell lines for transcriptomic profiling (Figures 1D and S1A). Pathway enrichment analysis was performed on the differentially expressed genes to identify cellular pathways that were regulated by MTHFD2. Surprisingly, the top metabolic pathway impacted by MTHFD2 loss was oxidative phosphorylation (henceforth abbreviated as OXPHOS; Figures 1D and S1B). We corroborated this result with tandem mass tag proteomics as well, noting dysregulation of multiple metabolic genes, including NADH: ubiquinone oxidoreductase core subunit S8 (NDUFS8), a core subunit of mitochondrial Complex I (Figure 1E). Many helicase genes were also differentially impacted, in agreement with an earlier report of MTHFD2 having a role in the regulation of RNA metabolism^9^. We further validated these results by performing a mitochondrial stress test with inhibitors of the mitochondrial electron transport chain and observed higher oxidative capacities in the MTHFD2-deficient cells relative to wild-type cells (Figures 1F and S1C). Increased oxidative capacities in MTHFD2-deficient cells was somewhat unanticipated, as the reduction in formate levels would have impacted mitochondrial translation^1^ and mitochondrial electron transport chain complex stoichiometry^13^. Inspection of the transcriptomic dataset revealed global downregulation of mitochondrial ribosome gene transcripts while mitofusin-1 (MFN1) showed significant up-regulation in MTHFD2-deficient cells. Interestingly, catalase (CAT), an enzyme that mediates peroxide detoxification, was also concomitantly upregulated (Figure 1E), presumably to reduce oxidative damage caused by increased OXPHOS. Together, these data suggest that MTHFD2 deficiency promotes mitochondrial consolidation, thereby enhancing oxidative potential.

### MTHFD2 supports creatine metabolism in B lymphoblastoid cells

Metabolomic analysis was then undertaken as a final confirmation of inferences from the gene expression datasets. As expected, serine accumulated in the cells that were deficient for MTHFD2 (Figure 2A); loss of flux through the mitochondrial pathway and subsequent reversal prevents rapid consumption of serine that causes buildup within the cell. Very few metabolites were differentially expressed between wild-type and MTHFD2-deficient cells. Interestingly, O-phosphoethanolamine (O-PEA) was the only other metabolite that was upregulated in MTHFD2-deficient cells. Oxaloacetate (OAA), a substrate of the tricarboxylic acid cycle (TCA), was depleted in MTHFD2-deficient cells, which was expected, given the higher oxidative profile of the cells. Apart from OAA, most of the other metabolites that were downregulated in MTHFD2-deficient cells, namely creatine, carnosine and taurine, were involved in bioenergetics and derived from amino acid metabolic pathways.

**Figure 2.**
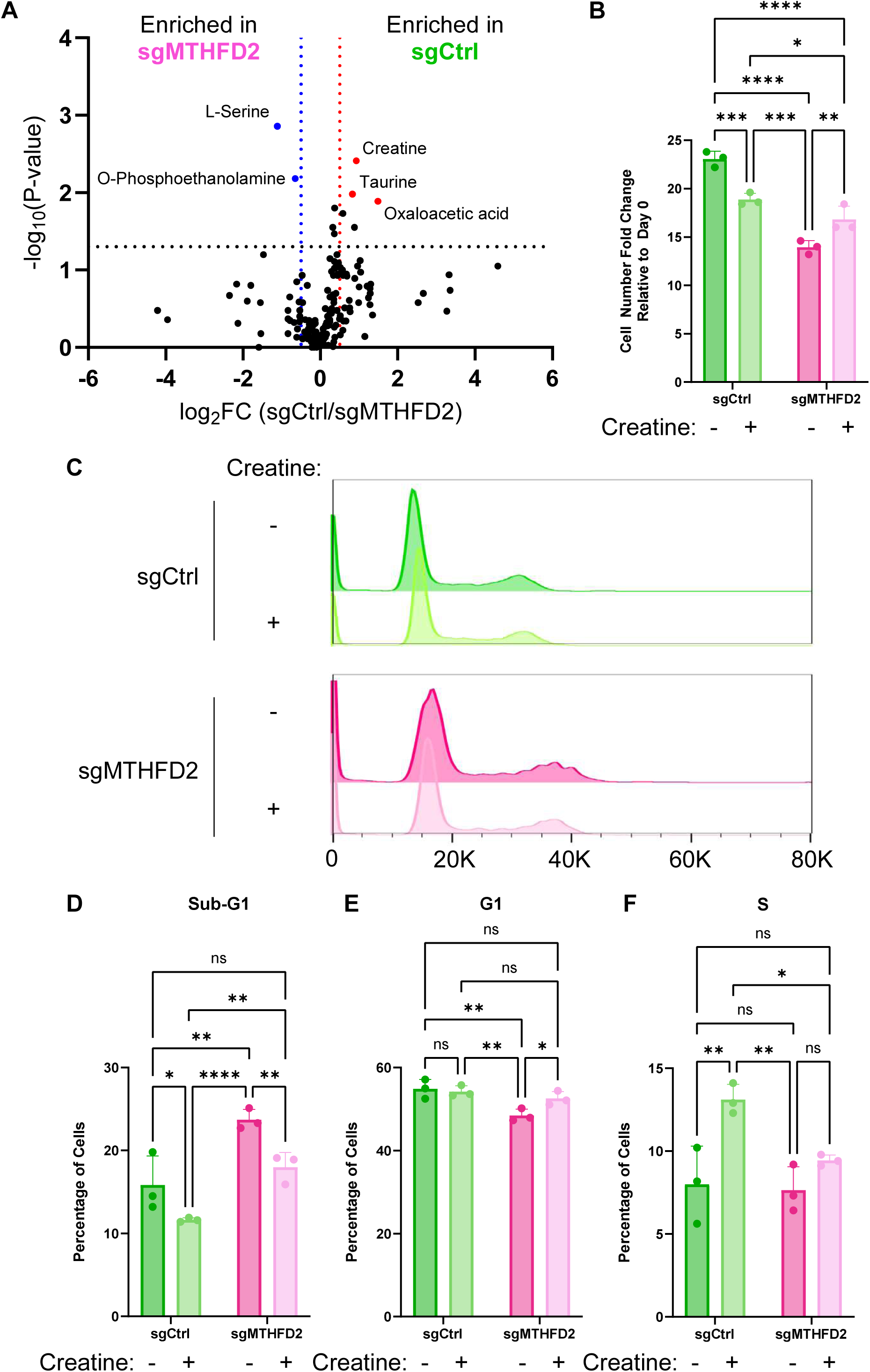
MTHFD2 supports creatine metabolism in B lymphoblastoid cells. (A) Volcano plot illustrating the differentially regulated metabolites in GM12878 sgCtrl and sgMTHFD2 cells. X-intercepts indicate fold-change cut-offs at X = -0.5 (blue) and X = 0.5 (red). The Y-intercept indicates the nominal P-value cut-off of 0.05. Blue dots indicate metabolites enriched in sgMTHFD2 cells whereas red dots indicate metabolites enriched in sgCtrl cells. (B) Growth analysis of GM12878 sgCtrl and sgMTHFD2 cells mock-treated or supplemented with creatine (10 mM). -, absence of supplement; +, presence of supplement. (C) Representative histograms for cell cycle analysis of GM12878 sgCtrl and sgMTHFD2 cells mock-treated or supplemented with creatine (10 mM). (D-F) Frequency plots of subpopulations from the cell cycle analysis shown in Panel C, namely (D) sub-G1, (E) G1 and (F) S phase cells. All data represent N = 3 biological replicates with errors bars indicative of SD. ns, not significant; *, p < 0.05; **, p < 0.01; ***, p < 0.001; ****, p < 0.0001 by Student’s t-test (Panel A) or ordinary two-way ANOVA with uncorrected Fisher’s LSD (Panels B, D-F). See also Figure S2.

We then asked the question whether any of these differentially expressed metabolites (DEMs) would rescue the viability defects induced by MTHFD2 loss. We systematically tested each of the notable DEMs i.e., creatine, taurine, OAA and O-PEA (Figures S2A-D), by supplementing them into culture at millimolar concentrations. Creatine supplementation was the only condition that rescued MTHFD2-deficient cell viability (Figures 2B and S2A). Taurine supplementation had no effect. Interestingly, supplementation with either OAA or O-PEA induced general decreases in cell viability through hitherto unknown mechanisms. Hereafter, we focused on creatine for subsequent experiments.

The mitigation of viability defects can be due to increased proliferation and/or reduced cell death. To determine how creatine affects these viability factors, we performed propidium iodide staining to assay cell cycle progression (Figure 2C) and CellTrace dye dilution assays to measure proliferation (Figure S2E). Creatine supplementation did not significantly alter the proliferative potential of the cells as indicated by similar CellTrace dye dilution profiles. However, significant changes were apparent in the propidium iodide staining. Creatine supplementation of MTHFD2-null cells reduced the sub-G1 population to wild-type levels, indicating anti-apoptotic effects (Figure 2D). G1 frequencies were also significantly elevated in MTHFD2-deficient cells supplemented with creatine and normalized to wild-type counterparts (Figure 2E), suggesting that creatine may also have a role in supporting non-proliferative cell functions. Intriguingly, creatine supplementation of wild-type cells but not MTHFD2-null cells had a significant impact on DNA synthesis (Figure 2F). G2/M progression was unaffected, regardless of MTHFD2 expression status or creatine exposure (Figure S2F).

### Creatine synthesis is a major sink of glycine generated by mitochondrial one-carbon metabolism

To determine the mechanism by which MTHFD2 regulates creatine levels, we performed U^13^C-serine isotope tracing experiments in wild-type and MTHFD2-deficient cells (Figure 3). As expected, MTHFD2 loss resulted in significant decreases in total glycine, including the (M+2) isotopologue; loss of the mitochondrial one-carbon branch results in backflow and flux reversal^10^ through serine hydroxymethyltransferase 1 (SHMT1). Additionally, formate levels were also decreased in MTHFD2-null cells as evidenced by lower (M+1) labeling in deoxythymidine monophosphate (dTMP), consistent with our expectations of strongly blunted mitochondrial 1C pathway flux.

**Figure 3.**
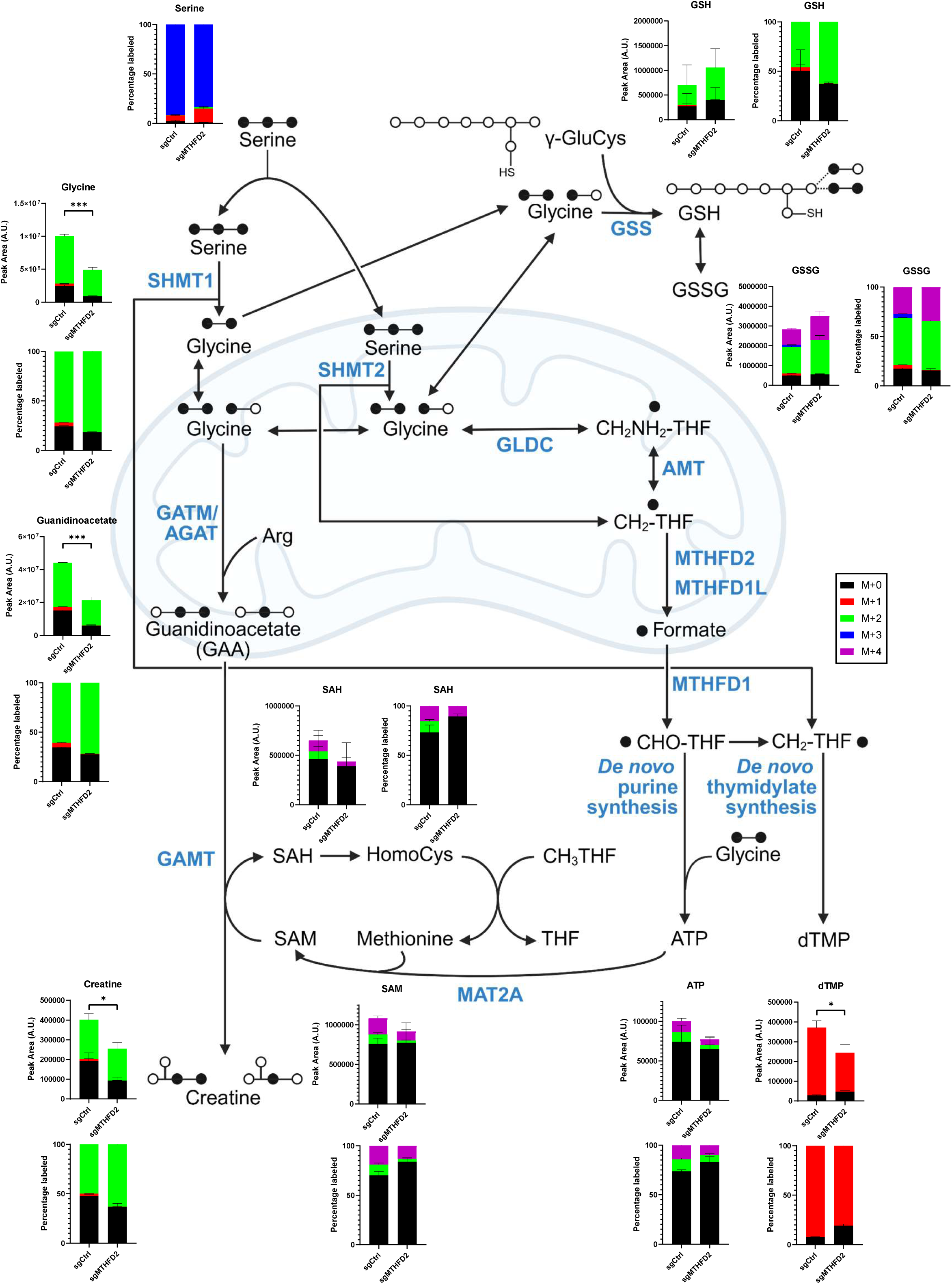
Creatine synthesis is a major sink of glycine generated by mitochondrial one-carbon metabolism. Plots of absolute and relative abundances of ^13^C-containing isotopologues for various metabolites across the indicated pathways in GM12878 sgCtrl and sgMTHFD2 cells after labeling with U^13^C-serine. Enzymes and metabolites are indicated in blue and black, respectively. Color coding for the isotopologues is shown as an inset. All data represent N = 3 biological replicates with errors bars indicative of SD. *, p < 0.05; ***, p < 0.001 by Student’s t-test. See also Figure S3.

To our surprise, we detected the occurrence of *de novo* creatine synthesis at steady state in our cell lines (Figure 3). This is unusual in that creatine synthesis is normally broken up into two spatially separated steps in humans; the first step of generating the pathway intermediate guanidinoacetate is mediated by AGAT/GATM and occurs at the kidney while the second step of converting guanidinoacetate into creatine is catalyzed by GAMT and completed in the liver.

We confirmed protein expression of GATM in LCL (Figure S3A) in agreement with a previously published proteomic dataset^2^. In contrast, when we assayed primary B cells activated by recombinant trimeric CD40 ligand (CD40L) and/or interleukin-4 (IL-4)^14^, GATM was not appreciably expressed (Figure S3A).

Isotopic analysis of the cell lines revealed that creatine synthesis was compromised in MTHFD2-deficient cells, as evidenced by the diminished levels of the pathway intermediate guanidinoacetate and creatine itself. Interestingly, other glycine-utilizing pathways were not significantly impacted; overall glutathione and adenosine triphosphate (ATP) levels were similar in wild-type and MTHFD2-deficient cells.

Close inspection of ATP isotopologues revealed that ^13^C atoms were incorporated either as ^13^C_2_-glycine to produce (M+2) ATP or as one ^13^C_2_-glycine plus two ^13^C_1_-formate to generate (M+4) ATP. Absolute levels and isotopologue distributions of S-adenosylhomocysteine (SAH) and S-adenosylmethionine (SAM) closely reflected that of ATP, with no heavier isotopologues observed. Further, while it is theoretically possible to convert SHMT-derived (M+1) 5,10-methylene-THF into (M+1) 5-methyl-THF for SAH re-methylation, the apparent absence of (M+1), (M+3) and (M+5) SAM indicates that LCLs do not engage in such an activity. Taken together, our results confirm that homocysteine re-methylation does not draw on one-carbon units derived from serine catabolism under methionine-replete conditions.

One intriguing observation was the appearance of (M+1) glycine in wild-type cells which was abolished in MTHFD2-deficient cells. This suggests that MTHFD2 has a role in driving reverse glycine cleavage system (rGCS) activity to augment mitochondrial glycine pools; glycine decarboxylase (GLDC) somehow mediates the abstraction of the (M+1) aminomethyl moiety bound to GCS H-protein (GCSH) and incorporate (M+0) bicarbonate to produce (M+1) glycine. Protein expression of GCSH, DLD and GLDC was observed in LCLs (Figure S3B). Intriguingly, AMT was the sole GCS subunit that was not detected by immunoblotting (Figure S3B). These observations are concordant with our previous proteomic study of LCLs as well as our analysis of public LCL transcriptomes^15^ (Figure S3C). This pattern of gene expression is compatible with rGCS activity; without AMT, forward GCS flux cannot proceed. We attempted to demonstrate labeling of cellular glycine with ^13^C-bicarbonate and regular U^12^C-serine in the media. Oddly, no ^13^C atoms were incorporated into cellular glycine or even into other metabolites, such as purines and pyrimidines, where bicarbonate is typically used for *de novo* syntheses. This is in spite of the fact that a bicarbonate transporter, SLC4A7, has been shown to be expressed in LCLs^2^.

Hence, it appears that the rGCS reaction in LCL may depend exclusively on endogenously produced bicarbonate. Cellular usage of (M+1) glycine is peculiarly asymmetric; its signal was detected in glutathione but not in ATP, despite both of their synthesis pathways being cytosolic. In other words, glutathione synthesis is dependent on glycine generated from two sources, namely SHMT-catalyzed reactions and rGCS activity. In contrast, purine synthesis draws exclusively on SHMT-derived glycine molecules, in agreement with a previous report showing close association between SHMT and the putative purinosome^16^.

Cellular metabolism often depends on the relative balance of metabolite pairs e.g. phospho-Cr/Cr (PCr/Cr), GSH/GSSG and SAM/SAH. Many of these metabolite pairs depend on glycine as a basic building block. Hence, we investigated whether their relative balances were impacted by MTHFD2 ablation. Overall and ^13^C-labeled GSH/GSSG and SAM/SAH ratios were not significantly altered whether MTHFD2 was expressed, or creatine was given as a supplement (Figures S3D-G). Interestingly, neither MTHFD2 loss nor Cr supplementation caused a significant change in overall PCr/Cr ratios, even though ^13^C-labeled PCr/Cr ratios strongly decreased with Cr supplementation (Figures S3H-I). The latter is expected given that Cr blocks its own synthesis in a negative feedback loop^17,18^.

To determine the role of virus- and/or host cell-encoded transcription factors in regulating creatine synthesis, we analyzed publicly available chromatin immunoprecipitation sequencing (ChIP-seq) datasets of GM12878 LCL^19^ (Figure S3J). At the *GATM* promoter, we observed the accumulation of EBV-encoded nuclear antigens, namely EBNA-2, EBNA-LP, EBNA-3A and EBNA-3C. In addition, nuclear factor kappa B (NF-κB) subunits were also enriched together with the EBNAs, suggesting that EBV-encoded latent membrane protein 1 (LMP-1), a known activator of NF-κB signaling^20,21^, may also function to augment GATM expression.

### Creatine synthesis is an endogenous tumor suppressor pathway hypostatic to mitochondrial one-carbon metabolism

Given that creatine supplementation rescued MTHFD2 knockout cell viability and that creatine levels were strongly diminished in the absence of MTHFD2, we hypothesized that ablating GATM would result in synthetic lethality against an MTHFD2-deficient background. To address this question, we performed dual CRISPR/Cas9 targeting at the *MTHFD2* and *GATM* gene loci to generate double knockouts (Figure 4A). Surprisingly, *GATM* deletion enhanced cell viability regardless of MTHFD2 expression status. In fact, GATM loss rescued MTHFD2 knockout-induced cell viability defects by significantly mitigating cellular apoptosis (Figure 4B). Consistent with that, millimolar supplementation of creatine, a known negative feedback regulator of GATM that had generated a rescue effect in MTHFD2-deficient cells (Figure 2), led to a significant reduction of intracellular guanidinoacetate (Figure S4A).

**Figure 4.**
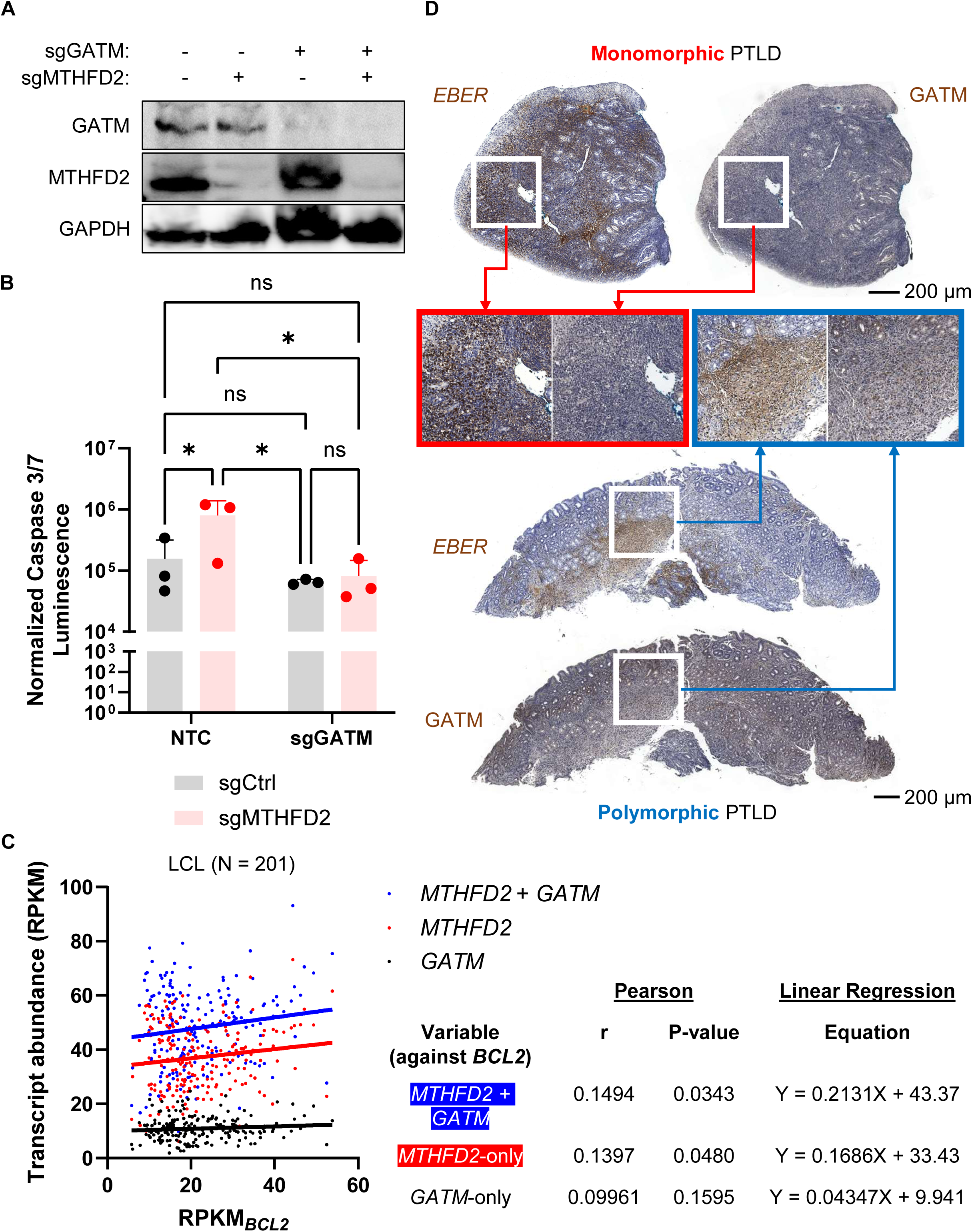
Creatine synthesis is an endogenous tumour suppressor pathway hypostatic to mitochondrial one-carbon metabolism. (A) Representative immunoblot (N = 2) of lysates from GM12878 cells singly or doubly ablated for MTHFD2 and GATM protein expression. GAPDH was used as housekeeping control. (B) Luminescence-based measurement of caspase 3/7 activity in GM12878 cells edited at the indicated gene loci, with sgCtrl or non-targeting control (NTC) sgRNA. Note that sgCtrl is the matched control to sgMTHFD2, encoded on a puromycin resistance marker-containing construct while NTC is the matched control to sgGATM, encoded on a hygromycin resistance marker-containing construct. (C) Correlation analysis of selected genes in public LCL transcriptomes (N = 201). For statistical testing, Pearson correlation coefficients were computed with the exact P-values indicated in the embedded table. (D) Representative scans of patient-derived gastrointestinal tissue sections showing *in situ* hybridization of nucleic acid probes to *EBER* and immunohistochemical staining of GATM protein. Scale bars (200 µm) are indicated. Inset frames show magnified views of the stained sections. N = 2 per sample from patients P2 and P6. Quantitative data represent N = 3 biological replicates with errors bars indicative of SD unless otherwise stated. ns, not significant; *, p < 0.05 by ordinary two-way ANOVA with uncorrected Fisher’s LSD (Panel B). See also Figure S4.

Creatine is eliminated from cells by the irreversible conversion of PCr into creatinine and subsequent export through transporters of the SLC22A and SLC47A families. Interestingly, creatinine was exclusively detected within cells and not in the culture supernatants. Consistent with this observation, members of the SLC22A and SLC47A families were not detected in LCL proteomic datasets^2^. Furthermore, MTHFD2 loss did not significantly alter intracellular creatinine levels, even though intracellular creatine levels were drastically diminished (Figures 3 and S4B). This suggests that flux through the creatine phosphorylation step was unaffected to maintain relatively steady levels of phosphocreatine. The net result of this is an increased PCr/Cr ratio, which is consistent with the bioenergetic patterns described earlier (Figures 1 and S3H).

Based on the working hypothesis that creatine synthesis has tumor suppressive effects, we considered whether GATM expression would influence PTLD progression and outcomes. To our knowledge, no database linking gene expression to clinical outcome exists for PTLD worldwide.

We made use of BCL2 as a proxy marker for disease severity as *BCL2* overexpression has been linked to worse prognoses in PTLD patients^22^. In our analysis of public LCL transcriptomes, *GATM* expression alone did not correlate with *BCL2* transcript abundance. However, when considered together with *MTHFD2* levels, improved correlation against *BCL2* was observed (Figure 4C). Next, we made use of TCGA patient-derived datasets^23^ for further testing of the hypothesis. As expected, high *GATM* levels were shown to be a positive prognostic factor in renal clear cell carcinoma^24^. We extended our investigation to *GAMT*, the other gene in the creatine synthesis pathway. Although GAMT detection was difficult to achieve with proteomics and antibody-based assays in our previous experiments, the TCGA analysis showed it to be a potentially positive prognostic factor in renal clear cell carcinoma as well^25^. Further, *GAMT* expression in LCLs was significantly and negatively correlated with a few PTLD-driving factors, such as the anti-apoptotic factor *CFLAR* (Figure S4C) and the immunosuppressive cytokine *IL-10* (Figure S4D).

To determine whether disease-driving cells in PTLD participate in *de novo* creatine synthesis *in vivo*, we performed *in situ* hybridization and immunohistochemical staining of tissue samples from monomorphic PTLD (more severe) and polymorphic PTLD (less severe) (Figure 4D).

GATM protein was appreciably detected in PTLD tissue regardless of severity. Intriguingly, *EBER1* signals appeared inversely correlated with GATM signals, consistent with active viral regulation of *GATM* gene expression (Figure S3J) and our analysis of public LCL transcriptomes (Figure S4E). Taken together, our data point to creatine synthesis operating as a tumor suppressor pathway in PTLD.

## Discussion

Although MTHFD2 is universally critical in supporting the growth and proliferation of lymphocytes and cancer cells, its specific functions appear to be highly varied in different cell types. Further, given its overexpression in various oncogenic programs, its apparent dispensability, i.e., minimal disruption to overall cell fitness, appears somewhat discordant.

Importantly, attempts to rescue MTHFD2 loss with known downstream metabolites have not been successful^7^. Overall, apart from its enzymology, MTHFD2 remains an enigmatic target.

To demystify the roles of MTHFD2 in the broader context of cell metabolism, we made use of multiple high-dimensional datasets spanning the transcriptome, proteome and metabolome to determine the effects of MTHFD2 ablation in EBV-transformed LCL. To our surprise, MTHFD2 loss conferred a hyper-oxidative phenotype to cells, underpinned in part by upregulation of the key OXPHOS protein NDUFS8. NDUFS8 augmentation has been shown to be critical in increasing mitochondrial respiration^26^. Strikingly, MTHFD2 loss also caused a severe shortfall of creatine. Creatine supplementation of lymphoblasts increases spare respiratory capacity^27^.

Conversely, creatine depletion in MTHFD2-null cells likely decreases spare respiratory capacity. Hence, the net hyper-oxidative phenotype observed in MTHFD2-null cells may represent metabolic adaptation to overcome the effects of creatine depletion.

Previous work by others have shown that creatine synthesis in humans is split between two distinct organs^28–30^. Here, we show that concerted activity of GAMT and GATM within the same cell can occur under pathophysiological conditions as with LCLs as a model of PTLD. Why might LCLs upregulate creatine synthesis? Creatine synthesis may have evolved as a means of disposing of excess glycine under a unique set of conditions, namely MTHFD2 overexpression together with an absence of AMT. Although glycine is required by rapidly proliferating cells to support nucleotide and glutathione syntheses^2,31^, it is likely to be produced in excess, together with formate overflow^32^, in our cell system. When MTHFD2 is lost, glycine production is severely diminished, resulting in less flux through creatine synthesis. Instead, some of the residual glycine is diverted towards glutathione synthesis, as evidenced by the slight elevation in (M+2) GSH and (M+4) GSSG levels. Notwithstanding these observations, other sinks for glycine may also exist. Creatine supplementation as a rescue agent for MTHFD2-null cells may act as a feedback regulator on GATM, reducing creatine synthesis flux to free up glycine for other purposes.

As alluded to earlier, excess glycine is usually disposed of by cells through the GCS and be released in the form of carbon dioxide (incorporated as bicarbonate) and ammonia (usually as ammonium ions). However, reverse GCS activity has not been reported to occur in eukaryotic systems, even though it is a pathway used by microbes to produce glycine from simpler, inorganic molecules. *In vitro* reconstitution of the GCS in *Escherichia coli*^33^ has yielded important insights, namely that high levels of GLDC (also known as GCSP) relative to other GCS subunits is critical to the production of glycine via reverse GCS activity, and that the rate of glycine synthesis is very slow at a k_cat_ of <0.1 s^-1^. Our findings are consistent in that GLDC was found to be upregulated to a greater extent than other GCS subunits and that the (M+1) species was a minor isotopologue in total cellular glycine. Given the evolutionary conservation of GCS genes between bacteria and humans^34^, biochemical principles gleaned from the study of the bacterial GCS may be applicable to our understanding of the human GCS.

We posit the following model. MTHFD2 expression in LCLs produces high amounts of formate and glycine that permit anabolic metabolism, including nucleotide synthesis. High levels of glycine necessitate a robust system for disposal to prevent toxicity. At baseline, the GCS is not efficient enough to remove excess glycine, and hence, creatine synthesis takes on the role of glycine clearance. Although the mitochondrial glycine transporter SLC25A38 can play a role in removing glycine and indeed has a four-fold lower K_m_ (higher affinity) for glycine than GATM (SLC25A38: 0.75 mM; GATM: 2.5-3 mM), its V_max_ is ∼2.5-fold lower (SLC25A38: 170 nmol min^-1^ mg^-1^ protein; GATM: 440-500 nmol min^-1^ mg^-1^ protein)^35–37^. Considering the minimal changes in isotopologue contributions to cytosolic sinks of glycine i.e., ATP and glutathione, subsequent to MTHFD2 depletion, it is likely that mitochondrial export of glycine is already maximal. On the flip side, when MTHFD2 is deleted, both formate and glycine levels decrease, resulting in less nucleotides being produced, and lower levels of creatine being generated. With low creatine levels, GATM undergoes de-repression to become more active. This exacerbates the glycine shortfall, making the cell cycle defects even more pronounced. However, supplementation of creatine or GATM deletion in MTHFD2-deficient cells prevents catastrophic depletion of glycine, thereby rescuing the cell from death and cell cycle arrest.

Serum creatinine is commonly measured in transplant patients after surgery due to the use of nephrotoxic immunosuppressants such as tacrolimus; drug-induced nephrotoxicity reduces the kidney’s ability to perform effective filtration and clearance of creatinine, leading to detectable accumulation in the blood. Our findings suggest that the cellular drivers of PTLD can likely also contribute to this signal through endogenous production of creatine and subsequently, spontaneous hydrolysis of its phosphorylated form. Although we did not detect creatinine in the extracellular milieu, this may be attributed to potential reasons, such as poor *in vitro* expression of creatinine exporter proteins in LCLs. *In vivo*, appropriate tissue microenvironmental signals may stimulate the LCL-like cells to upregulate transporter proteins that can efficiently discharge creatinine into systemic circulation. Further, serum creatinine levels positively associate with graft failure in patients^38^. For sufficiently severe PTLD, creatinine produced by LCL-like cells may confound clinical interpretation of serum creatinine levels, presenting a diagnostic possibility that is overlooked i.e., a rise in serum creatinine not due to nephrotoxicity but rather, consequent to PTLD. This has potential effects on the assessment of drug-induced nephrotoxicity and clinical decisions to continue or cease immunosuppressant treatment. To obviate this issue, it is important to utilize complementary approaches to assess renal function and it reinforces the necessity of combinatorial monitoring of urinary metabolites and proteins in PTLD patients.

## Resource Availability Lead Contact

Requests for further information and resources should be directed to and will be fulfilled by the Lead Contact, Liang Wei Wang.

## Materials Availability

Plasmids and cell lines generated in this study will be made available on request. However, payment and/or a completed materials transfer agreement might be required if there is potential for commercial application.

## Data and Code Availability

Transcriptomic data have been deposited at GEO with the dataset identifier GSE296284. Mass spectrometry proteomics data have been deposited to the ProteomeXchange Consortium via the PRIDE partner repository with the dataset identifier PXD063247. Any additional information required to re-analyze the data reported in this paper is available from the Lead Contact upon request.

## Acknowledgements

This work was funded by SIgN, Agency for Science, Technology and Research (A*STAR) via the Central Research Fund – Use-Inspired Basic Research [BMRC CRF (UIBR)] award and a Singapore Ministry of Health Open Fund – Young Investigator Research Grant (MOH-000545), both awarded to L.W.W. We acknowledge the assistance of the following entities: SIgN Immunomonitoring Platforms [funded by the National Research Foundation (NRF) Immunomonitoring Service Platform (ISO) Grant (NRF2017_SISFP09)], SIgN Immunogenomics, and PANOMIX Singapore Pte. Ltd.

## Author Contributions

Conceptualization, N.Y.T.L., L.W.W.; Methodology, N.Y.T.L., L.W.W.; Validation, N.Y.T.L., L.W.W.; Formal Analysis, N.Y.T.L., R.A.T., R.B., N.A., H.S., L.W.W.; Investigation, N.Y.T.L., T.M.Y., R.A.T., L.L., C.Y.C., D.W.Q.L., M.Y.L., J.A.P., H.S., L.W.W.; Resources, J.A.P., W.W., H.S., L.W.W.; Writing – Original Draft, N.Y.T.L., L.W.W.; Writing – Review & Editing, N.Y.T.L., L.W.W.; Visualization, N.Y.T.L., R.B., N.A., L.W.W.; Supervision, W.W., H.S., L.W.W.; Funding Acquisition, J.A.P., W.W., H.S., L.W.W.

## Declaration of Interests

The authors declare no competing interests.

## Supplemental Information

Document S1. Figures S1-S4 and Table S1.

## STAR Methods

### • Key Resources Table

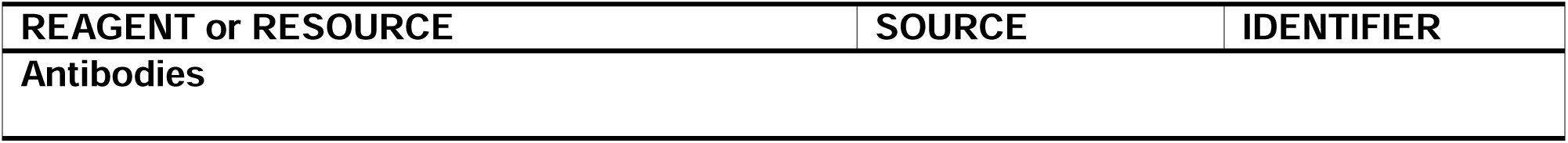
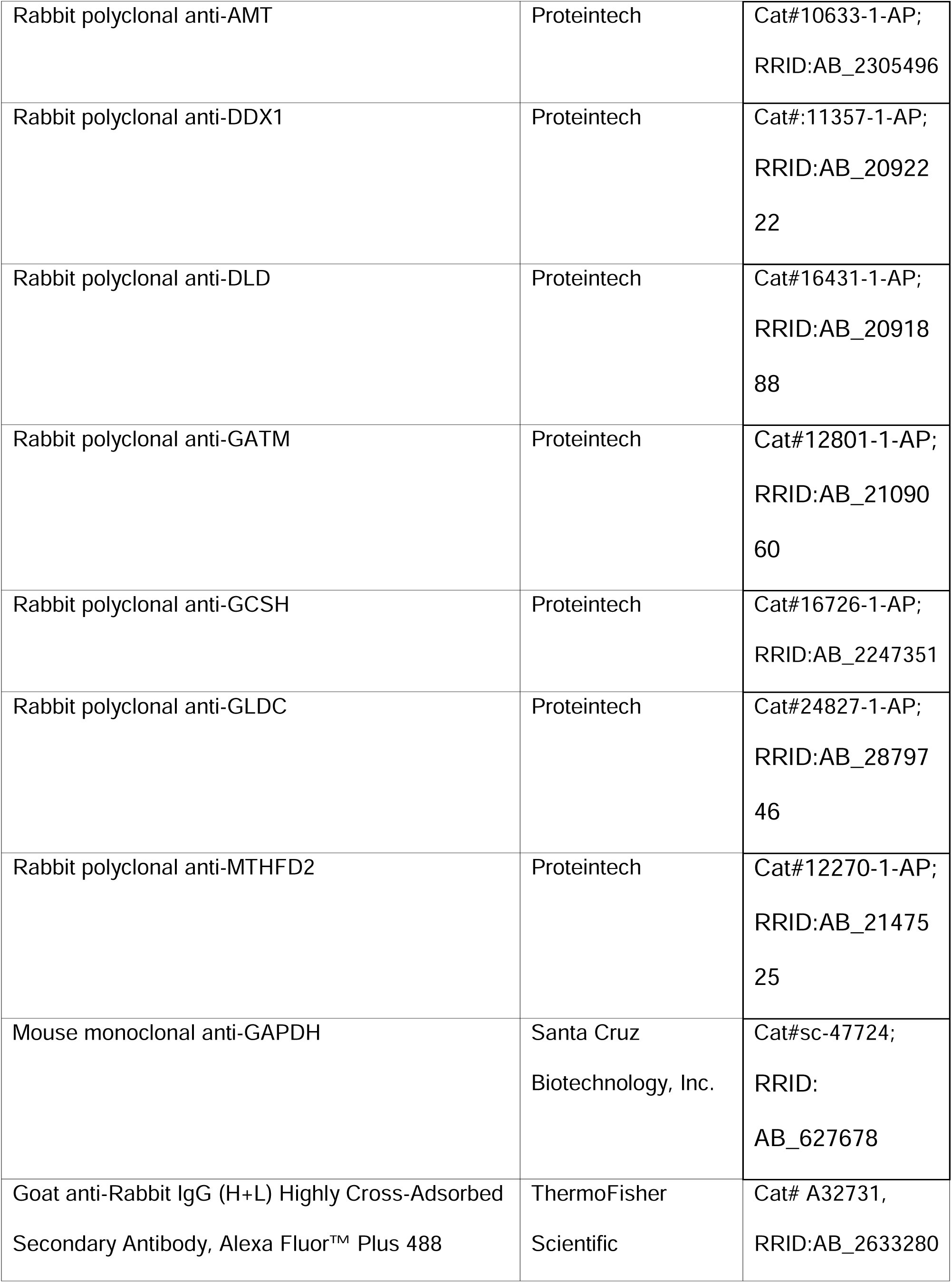
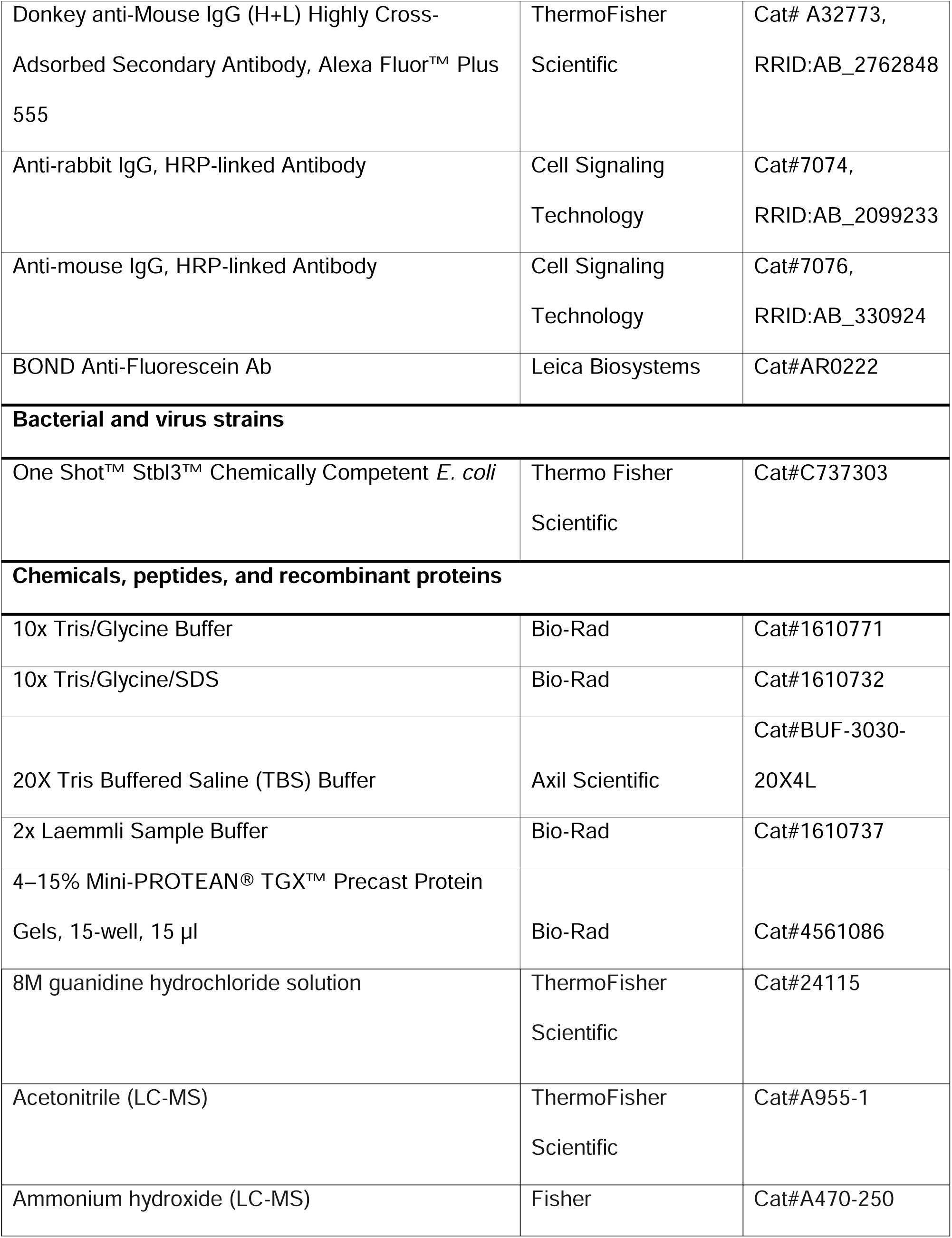
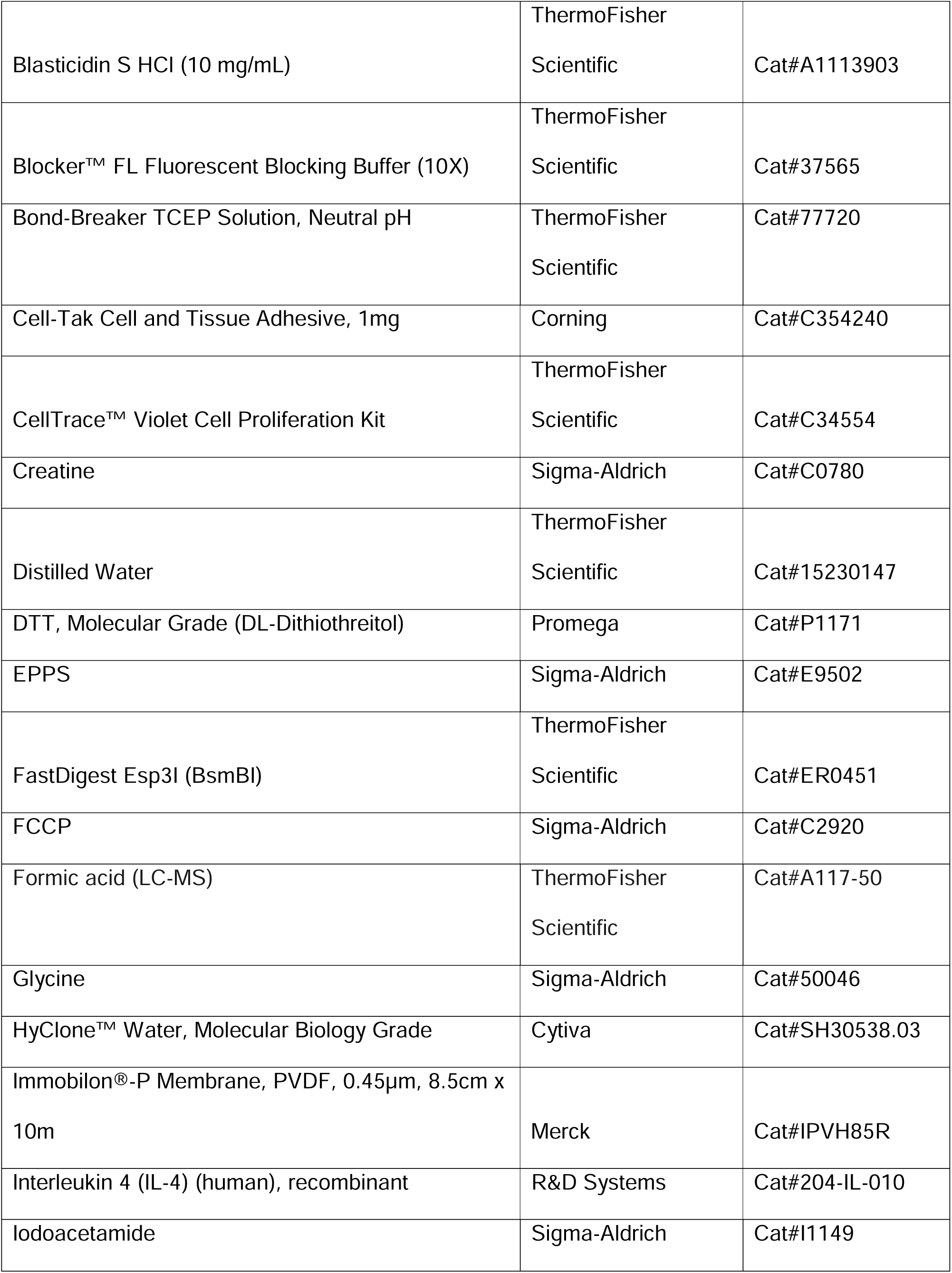
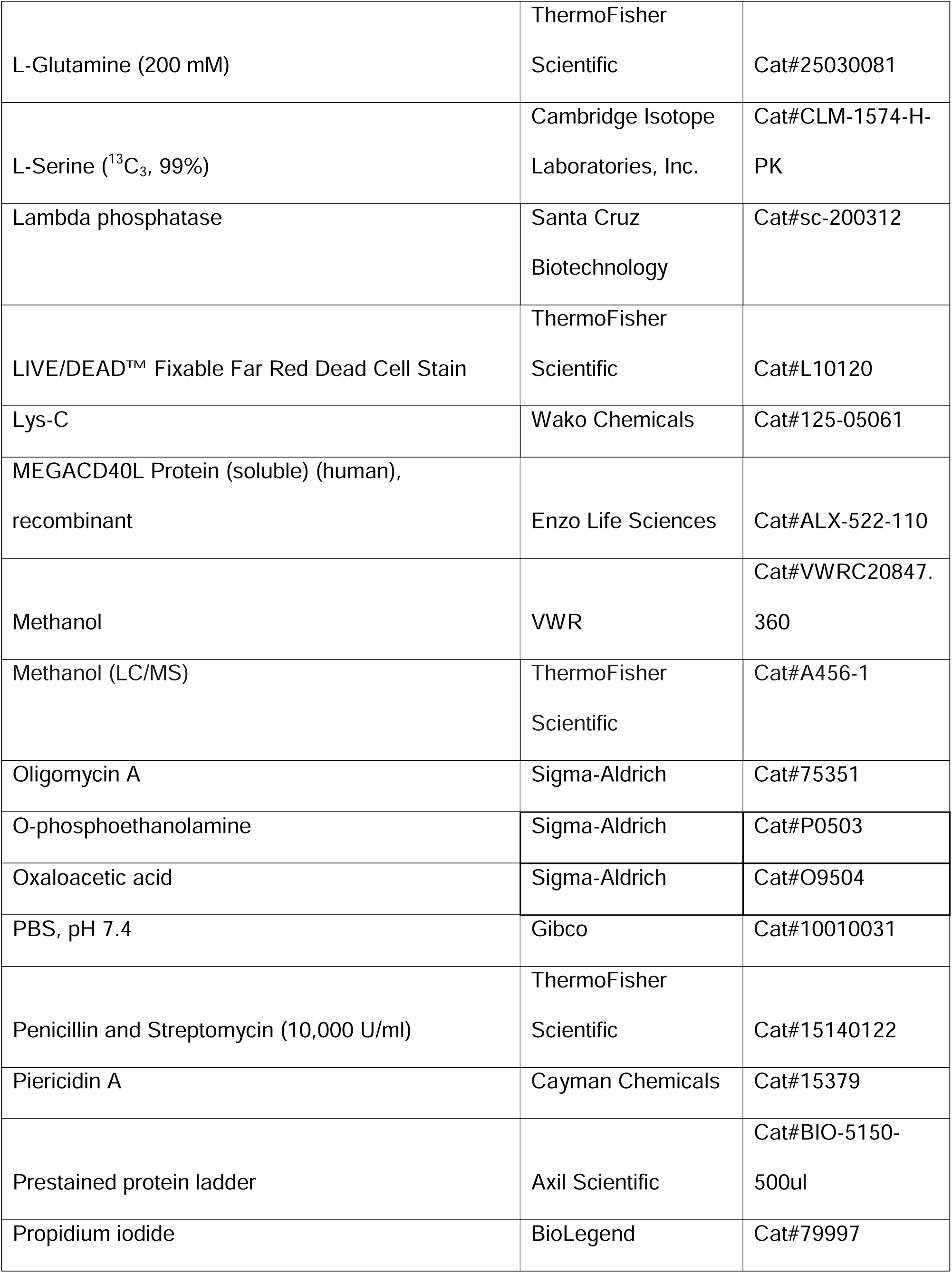
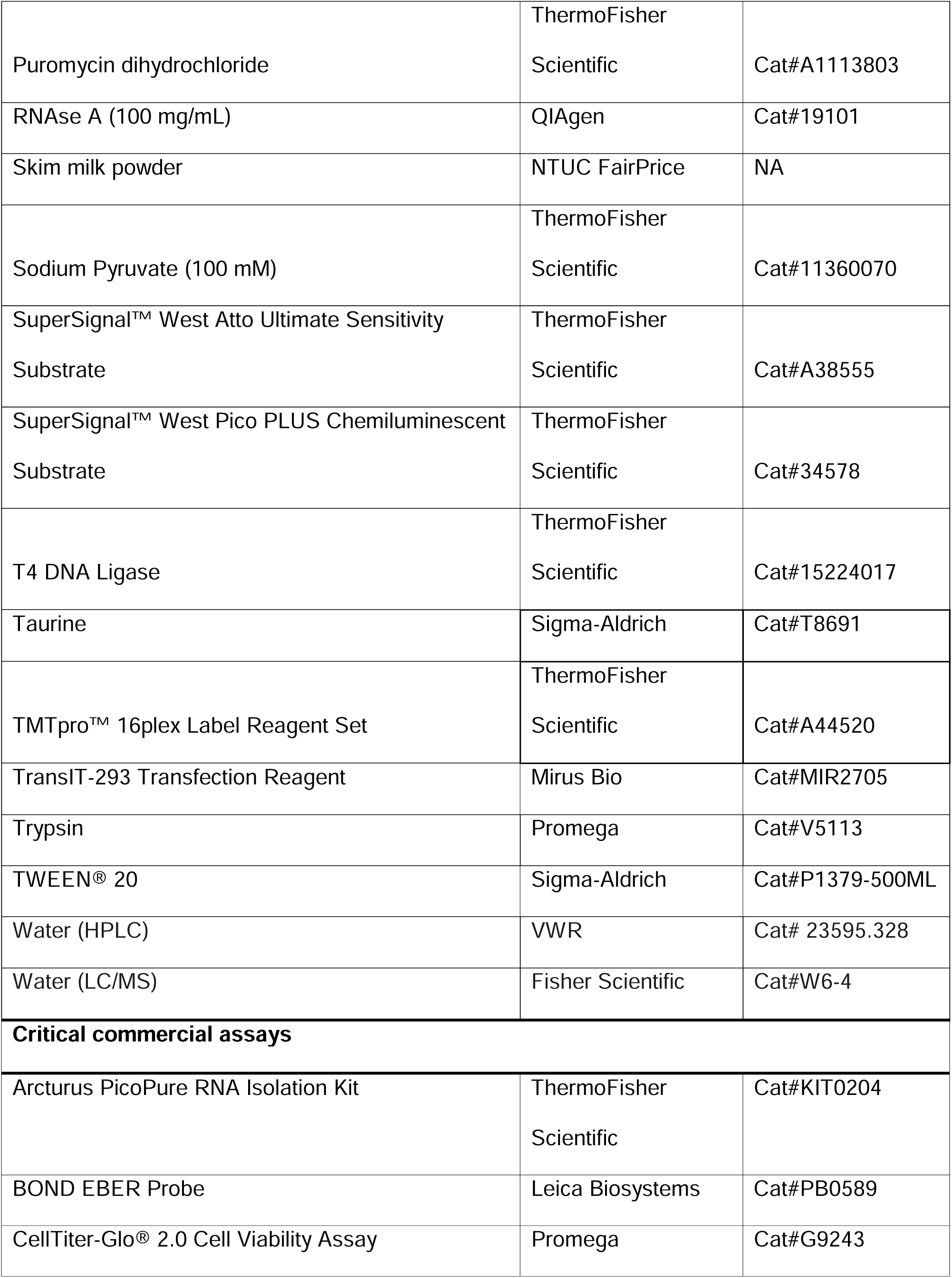
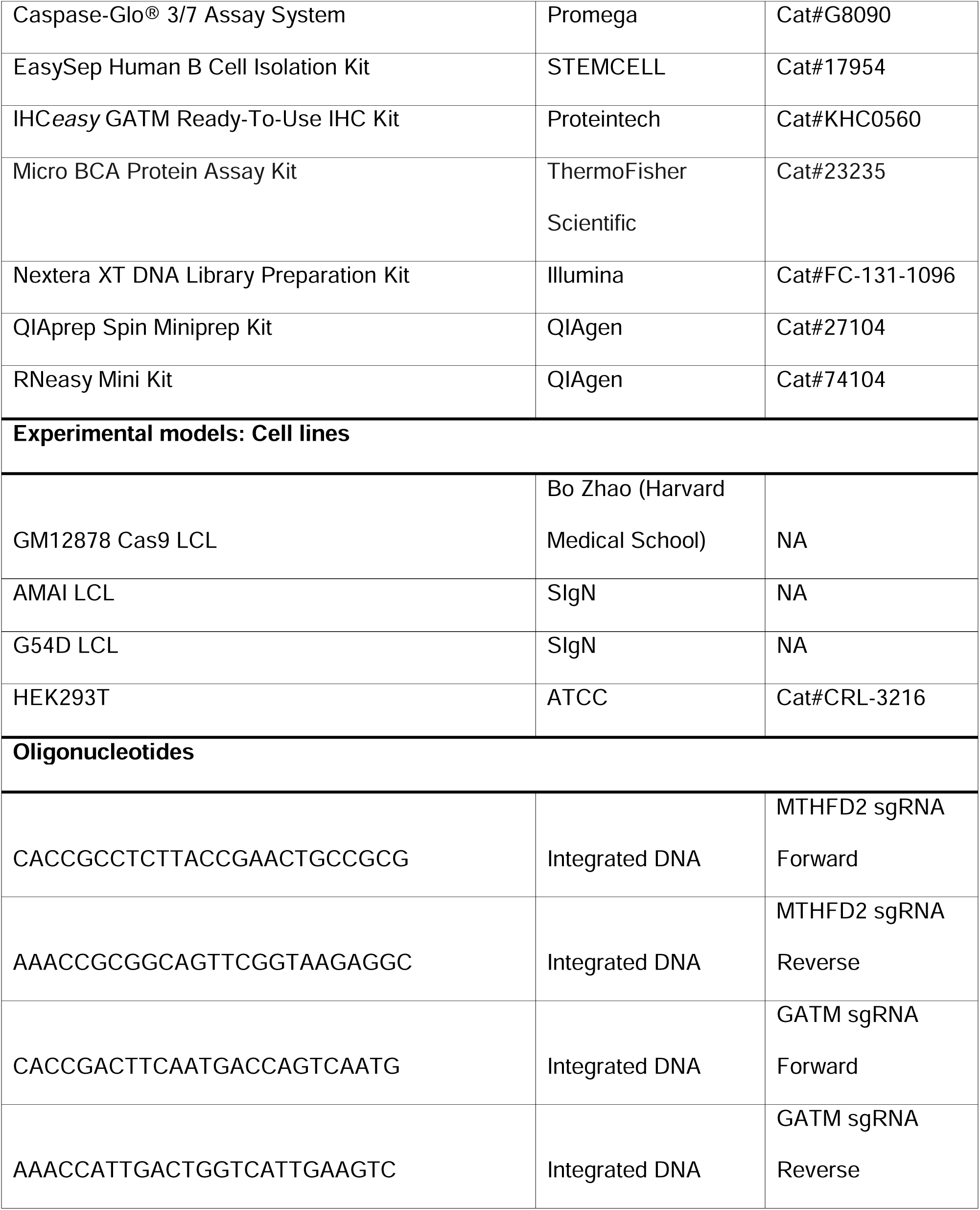
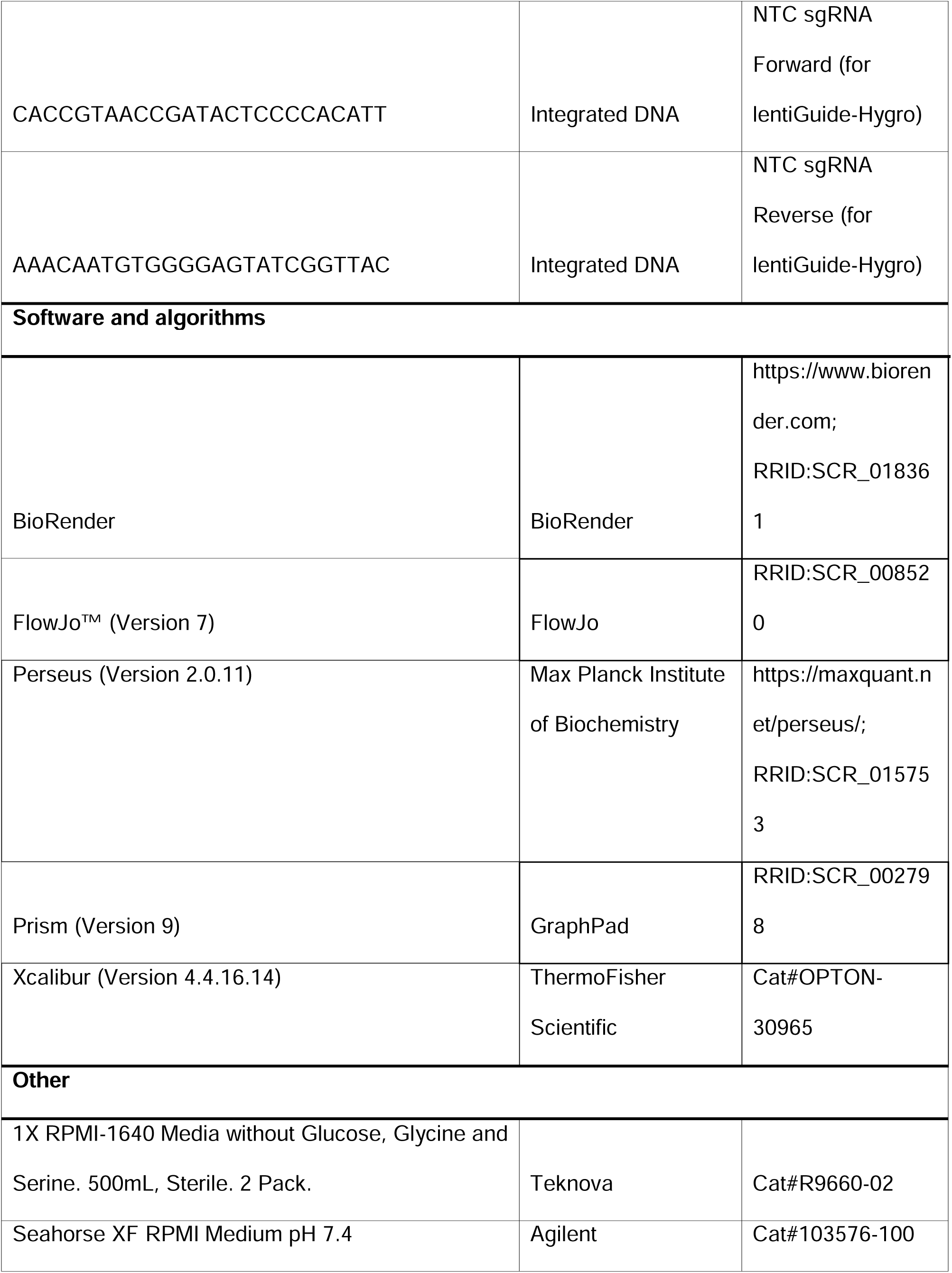
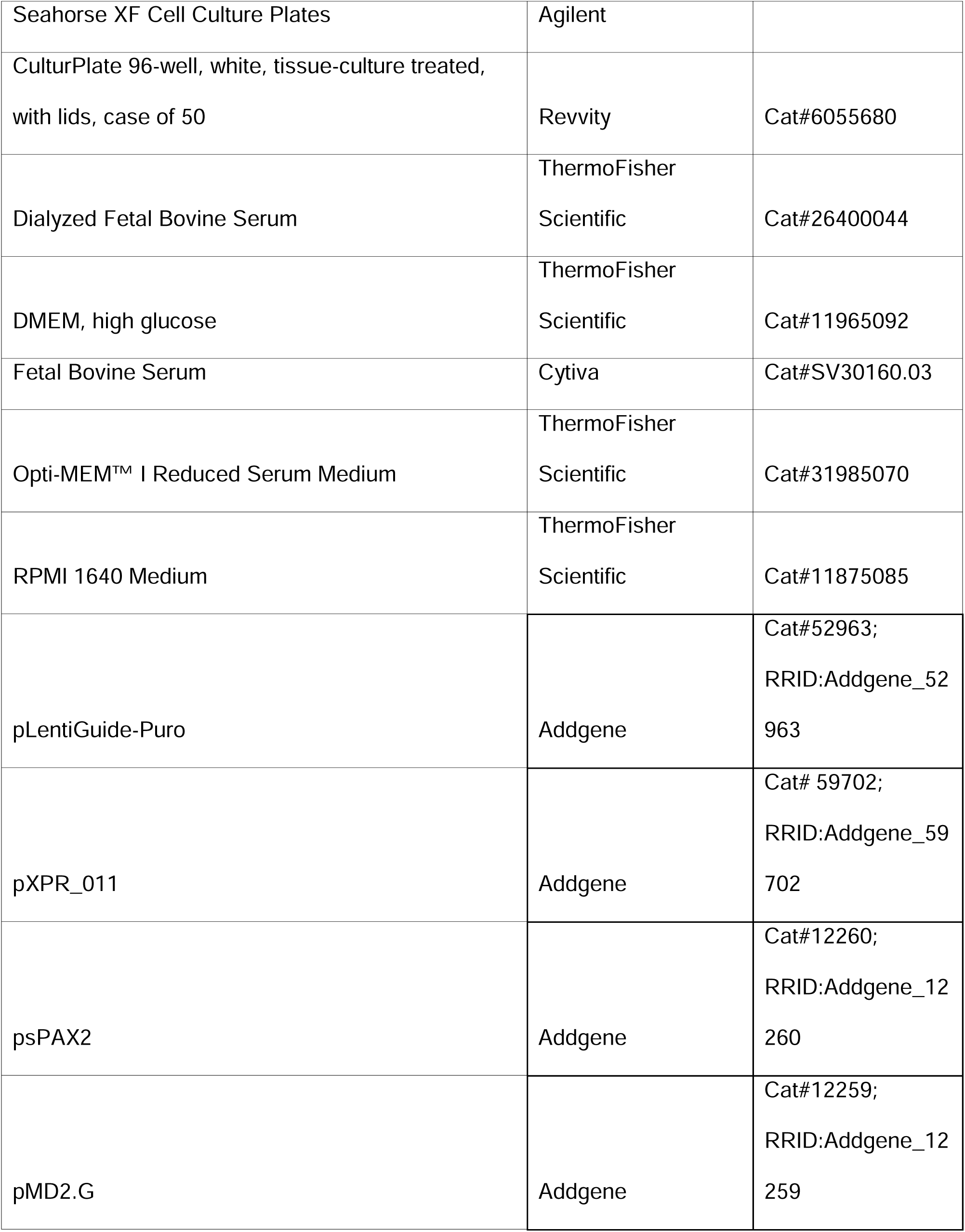
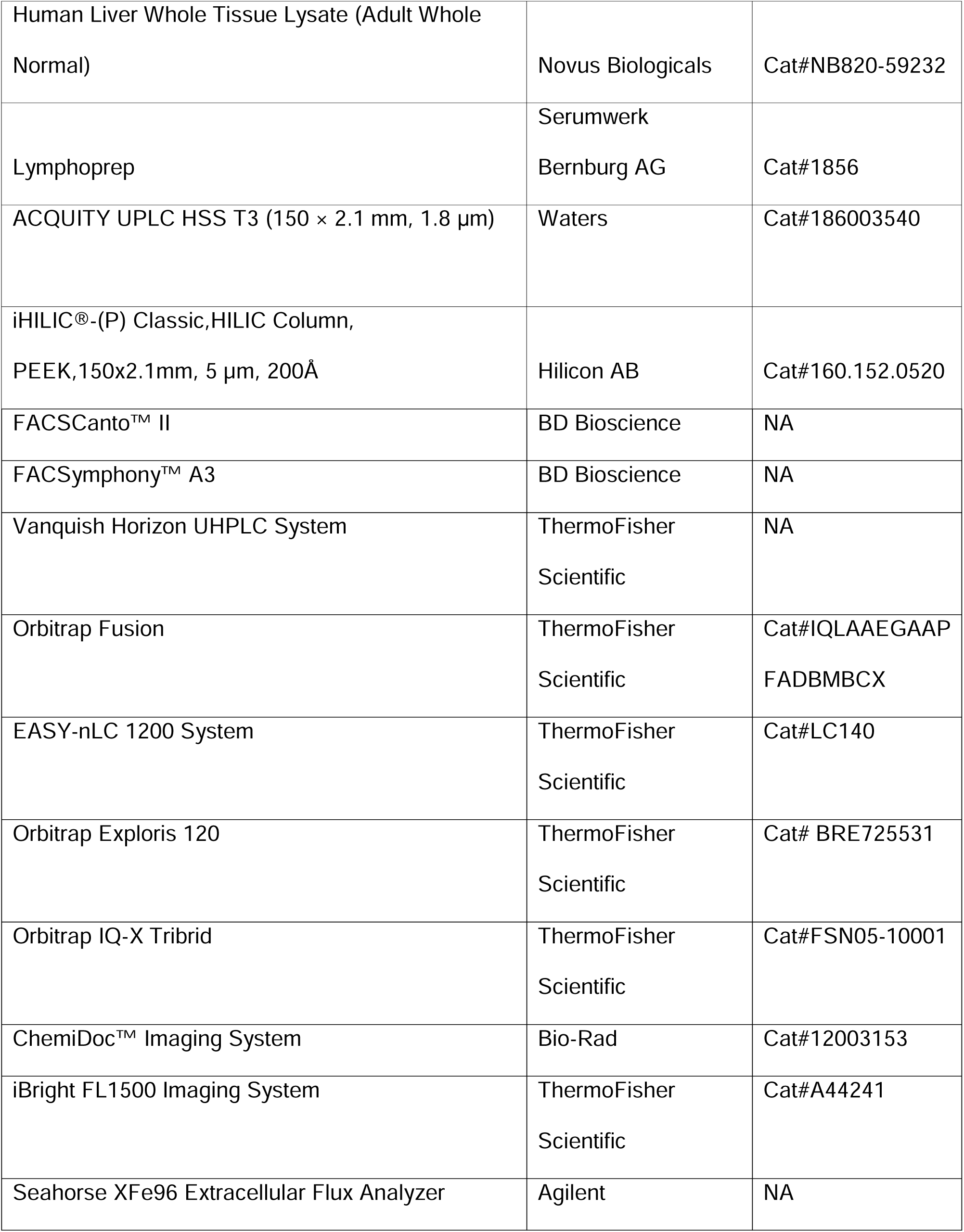

#### • Contact for Reagent and Resource Sharing

Further information and requests for resources and reagents may be directed to Liang Wei Wang (Lead Contact and Corresponding Author).

### o Experimental Model and Subject Details

#### • Culture of Established Cell Lines

HEK293T cells were cultured in DMEM with 10% heat-inactivated fetal bovine serum (FBS) and 1% Penicillin-Streptomycin. GM12878 lymphoblastoid cells (LCLs) originated from Coriell. Lentiviral transduction with the lentiCas9-Blast plasmid was utilized to generate GM12878 Cas9 cell lines, with transduced cells undergoing 2 weeks of selection with blasticidin at 40 μg/mL. Both GM12878 and GM12878 Cas9 cell lines were cultured in RPMI-1640 supplemented with 10% non-heat inactivated dialyzed FBS and 1% Penicillin-Streptomycin. For selection of post-lentiviral transduced cell lines stably expressing sgRNA, puromycin was used at 3.33 μg/mL. All cells were cultured in a humidified incubator at 37°C and at 5% CO_2_.

#### • Primary Human Tissue Samples

De-identified human blood tissue was collected in accordance with the approved IRB studies titled “Study of blood cell subsets and their products in models of infection, inflammation and immune regulation” (CIRB Ref: 2017/2806 and A*STAR IRB Ref. No. 2024-003). De-identified human blood tissue was also collected in accordance with and under the following project: HSA Residual Blood Samples for Research, project titled “Harnessing immune response for new therapies in transplantation and cancer” (HSA Study Ref. No. 201306-04 and A*STAR IRB Reference Number: 2024-133).

Peripheral blood mononuclear cells (PBMCs) were isolated from whole blood and cone blood samples by Lymphoprep centrifugation. Negative selection of B cells was performed with the EasySep Human B Cell Isolation Kit according to manufacturer’s instructions. Isolated B cells were enumerated and seeded in complete growth media (RPMI-1640 supplemented with 10% heat-inactivated FBS and 1% Penicillin-Streptomycin) at 1 million cells/mL. Stimulation with physiological agonists was performed as previously described^2,14^, using MEGACD40L and interleukin-4 (IL-4) at 50 ng/mL and 10 ng/mL, respectively. Cells were replenished with cytokines at 3 days and harvested after 7 days for lysate preparation.

#### • Pediatric PTLD Subjects

Anonymized histological specimens of EBV-positive B-cell post-transplant lymphoproliferative disease from pediatric patients who had previously undergone solid-organ transplantation were obtained from the Department of Pathology, National University Hospital, Singapore. For controls, tonsillectomy specimens with a histological diagnosis of reactive lymphoid hyperplasia were similarly obtained. As specimens and the accompanying data were anonymized, the study was reviewed by the National University Health System Research Office and certified as exempt from Institutional Review Board review (RNR2024-0011). Basic patient details are provided in Table S1.

### o Method Details

#### • CRISPR Editing in GM12878 Cas9 LCL

Single guide RNA (sgRNA) constructs were generated as previously described^12^. Sequences were obtained from the Brunello library^39^. MTHFD2 sgRNA (sgMTHFD2) oligonucleotides were synthesized and cloned into pLentiGuide-Puro and finally validated by Sanger sequencing.

LCL CRISPR editing was also performed as previously described^12^. Briefly, lentiviruses encoding sgRNAs were generated by transient transfection of 293T cells with packaging plasmids and either pLentiGuide-Puro sgMTHFD2 or pXPR_011. GM12878 cells stably expressing Cas9 were transduced with the lentiviruses and selected with 3 μg/mL puromycin for three days before replacement with antibiotic-free media. Single cell cloning by limiting dilution was performed to select for three clones each of GM12878 sgMTHFD2 knockouts and GM12878 pXPR_011 negative controls (sgCtrl).

#### • Growth Curve Analysis

Experiments were performed as previously described^2,40^. Briefly, GM12878 sgMTHFD2 and sgCtrl cells were seeded at 100,000 cells/mL at a volume of 1 mL of complete media (RPMI-1640 supplemented with 10% non-heat inactivated dialyzed FBS and 1% penicillin-streptomycin) in a 24-well plate. Cells were cultured for 7 days in total in 10 mM of the indicated metabolite supplement and sampled at regular intervals for live cell enumeration. At day 2, 4 and 7 post-seeding, cells were pelleted and resuspended in the same volume of complete media described earlier. Cells were stained with trypan blue and enumerated for live cells with a hemocytometer. At confluence, cells were split at a ratio of 1:4, where 250 µL of homogenous cell suspension was added to 750 µL of fresh media, maintaining total volume at 1 mL. Final cell counts were computed, considering the passage ratio. At each time point, cells were treated with fresh doses of metabolites at 10 mM after passaging.

#### • Mitochondrial Stress Test

Experiments were performed as previously described^2^. Briefly, Seahorse XF cell culture plates were coated with Cell-Tak solution to enable adhesion of GM12878 sgMTHFD2 and sgCtrl cells that were seeded at 200,000 cells per well. Complete phenol red-free XF RPMI-1640 media supplemented with 10% non-heat inactivated dialyzed FBS, glucose, glutamine and sodium pyruvate, was used as the culture medium over the assay period. Oxygen consumption rate was detected using the Seahorse XF96 sensor cartridges with data acquired via the Seahorse XF96 Extracellular Flux Analyzer. Mitochondrial poisons were used at the following concentrations – 3.5 µM oligomycin, 2 µM FCCP and 100 nM piericidin A.

#### • Transcriptome Extraction, Sequencing and Analysis

Total RNA was extracted from cells using the Arcturus PicoPure RNA Isolation kit according to the manufacturer’s protocol. The RNA samples were analyzed on Agilent Bioanalyser for quality assessment with RNA Integrity Numbers (RIN) ranging from 8.9 to 9.9. cDNA libraries were prepared using 2 ng of total RNA using the SMARTSeq v2 protocol^41^ with the following modifications: (a) addition of 20 µM TSO, and (b) use of 200 pg cDNA with 1/5 reaction of Illumina Nextera XT kit. The length distribution of the cDNA libraries was monitored using a DNA High Sensitivity Reagent Kit on the Perkin Elmer Labchip. Samples were subjected to 2x151 cycle sequencing on an Illumina HiSeq X or NovaSeq 6000 targeting at least 13 million read-pairs per sample.

Quality assessment of sequence reads in FASTQ files was done using FastQC^42^ and MultiQC^43^, with all samples passing QC. Paired-end (PE) reads were then aligned to the GRCh38 human genome using STAR^44^. The number of PE reads per sample ranged from 12.6-18.4 million, with a median of 15.6 million. Of the aligned reads, a median of 88.6% were uniquely mapped, with per-sample mapping rates ranging from 85.1%-87.6%. Gene counts were calculated using featureCounts^45^ based on Gencodev29 gene annotations^46^. Between 47.5% and 56.9% of mapped reads were assigned to genes, with a median of 50.5%. Log2 transformed counts per million mapped read (log2CPM) and log2 transformed reads per kilobase per million mapped reads (log2RPKM) values were computed using edgeR^47^. Genes with inter-quartile range (IQR) of log2CPM less than 0.5 across samples were excluded from subsequent differential expression analysis. edgeR was used to identify differentially expressed genes and Principal Component Analysis (PCA) on log2RPKM values was performed using the R function ‘prcomp’.

#### • LC–MS/MS Sample Preparation, Data Acquisition and Analysis for Proteomes

Cells were washed twice with PBS, and 150 µl of 6M guanidine/50 mM HEPES pH 8.5 lysis buffer was added. Samples were vortexed extensively then sonicated. Cell debris was removed by centrifuging at 13,000 x g for 10 min twice. Dithiothreitol (DTT) was added to a final concentration of 5 mM and samples were incubated for 20 min. Cysteines were alkylated with 15 mM iodoacetamide and incubated for 20 min at room temperature in the dark. Excess iodoacetamide was quenched with DTT for 15 min. Samples were diluted with 200 mM HEPES pH 8.5 to 1.5 M guanidine, followed by digestion at room temperature for 3 hours with LysC protease at a 1:100 protease-to protein ratio. Trypsin was then added at a 1:100 protease-to-protein ratio followed by overnight incubation at 37°C. The reaction was quenched with 1% formic acid. Samples were spun at 21,000 x g for 10 min to remove debris and undigested protein and then subjected to C18 solid-phase extraction (Sep-Pak, Waters) and vacuum-centrifuged to near-dryness. In preparation for TMT labelling, desalted peptides were dissolved in 200 mM HEPES pH 8.5. Peptide concentrations were measured by Micro BCA assay, and 50 mg of peptide was labelled with TMT reagent. TMT reagents (0.8 mg) were dissolved in 43 μl anhydrous acetonitrile and 5 μl was added to each peptide sample at a final acetonitrile concentration of 30% (v/v). Samples were labeled as follows: MTHFD2 sgRNA Replicate 1 (TMT 126), MTHFD2 sgRNA Replicate 2 (TMT 127N), MTHFD2 sgRNA Replicate 3 (TMT 127C), pXPR_011 Replicate 1 (TMT 130N), pXPR_011 Replicate 2 (TMT 130C), pXPR_011 Replicate 3 (TMT 131). Following incubation at room temperature for 1 hour, the reaction was quenched with hydroxylamine to a final concentration of 0.5% (v/v). TMT-labeled samples were combined at a 1: 1: 1: 1: 1: 1 ratio. The sample was vacuum-centrifuged to near dryness and subjected to C18 solid-phase extraction. Offline high pH reversed-phase fractionation of peptides was performed and combined as described previously^48^.

Samples were analyzed with Orbitrap Fusion with FAIMSPro in data dependent mode. Fractionated samples were reconstituted in solvent A (0.1% formic acid). Peptides were separated using a 90-min linear gradient up to 80% acetonitrile in 0.1% formic acid. For acquisition methods, FAIMS switching was set to 1 second for -40, -60 and -80 compensation voltage (CV). The spray voltage was set to 2.6kV and temperature of the ion transfer tube was set to 300◦C. MS1 scans were acquired at a resolution of 120k within the m/z range of 375– 1500 and automated gain control (AGC) target was set to 4e5. Multiply charged precursor ions starting from m/z 120 were selected for further fragmentation. Collison induced dissociation (CID) was performed with 35% normalized collision energy (NCE), and with dynamic exclusion of 60 s and 0.7 m/z isolation window MS2 scans were acquired within mass range of 400-1200 and the AGC target was set to 2e5. MS2 isolation window m/z was set at 2 for selection of ions for MS3. Higher energy collisional dissociation (HCD) was performed with 65% NCE, and with dynamic exclusion of 60 s and 1.3 m/z isolation window. MS3 scans were acquired at a resolution of 50k within the m/z range of 100–500. Automated gain control (AGC) target was set to 1.5e5.

MS data was acquired using ThermoScientific Xcalibur version 4.4.16.14. Raw files were searched against the human UniProt database (April 2024 with 204,093 SwissProt reviewed entries) including common contaminants using MaxQuant software version 2.5.2.0 with the integrated Andromeda search engine. For all files, standard parameter settings were used with Reporter MS3 for TMT11plex isobaric labels as the quantitative method with appended correction factors. Cysteine carbamidomethylation was set as fixed modification. Protein N-terminal acetylation and methionine oxidation were allowed as variable modifications. Digestion enzyme was set as Trypsin/P, allowing up to two missed cleavages. A false discovery rate (FDR) of 1% was used for peptide and protein identification.

Data was analyzed using Perseus software version 2.0.11. Corrected reporter intensities of proteins were log_2_ transformed. For further analysis, proteins quantified in at least two out of three replicates in at least one experimental condition were retained, and missing values were imputed based on normal distribution. To filter for significant changes between experimental conditions, two-sample paired Student’s t test were performed. FDR-corrected p values (q-values) were calculated from 250 randomizations and considered significant if they were 0.05 or less.

#### • Flow Cytometry

CellTrace Violet (CTV) staining was used to measure cell proliferation. Cells were initially seeded at 300,000 cells/mL and stained with CTV at a final concentration of 2 µM. Excess dye was subsequently quenched using neat ice-cold non-heat inactivated dialyzed FBS and washed with ice-cold complete media. Cells were then cultured in pre-warmed complete media as described earlier. As a control, a subset of stained cells was taken for flow cytometry analysis on day 0. Prior to flow cytometry analysis, the CTV-stained cells were washed once with room-temperature PBS and incubated with LIVE/DEAD™ Fixable Far Red Dead Cell Stain (reconstituted with DMSO as per manufacturer’s instructions) at a dilution of 1:500 for 10 minutes at room temperature, in the dark. Cells were then pelleted and resuspended in 400 μL of PBS, transferred into flow cytometry-compatible 5 mL tubes and processed immediately with the FACSCanto™ II flow cytometer. CTV-stained cells were left to culture over three days before the aforementioned analysis was carried out as well.

Propidium iodide (PI) staining was used to assess cell viability and the proportion of cells under each cell cycle state. Prior to staining, cells were treated over seven days with either creatine supplementation (10 mM) or distilled water (solvent control). Cells were harvested and washed twice with ice-cold PBS and pelleted before being fixed in 1 mL of ice-cold 70% ethanol and washed twice thereafter. Cell pellets were then treated with RNase A at 10 µg/mL, stained with PI at 2 µg/mL in a final volume of 250 µL and incubated at 4°C in the dark. The suspension was then transferred into 5 mL tubes and processed with the FACSymphony™ A3 flow cytometer.

#### • Untargeted Metabolomics

GM12878 sgMTHFD2 and sgCtrl cells were washed with PBS twice, aspirated and flash-frozen as cell pellets for conveyance to PANOMIX Singapore for metabolomic sample processing. A volume of 1 mL of ice-cold acetonitrile (ACN): methanol: water solution (2:2:1, v/v/v) was added to cell pellets and the lysate was vortexed for 30 seconds before centrifugation for 10 min at 12,000 rpm and 4°C. Clarified supernatants were concentrated and dried. A volume of 300 μL ice-cold acetonitrile: 2-amino-3-(2-chloro-phenyl)-propionic acid (4 ppm) solution prepared with 0.1% formic acid (1:9, v/v) was added to lyophilized samples for redissolution. Samples were filtered with 0.22 μm membranes and transfer into LC-MS compatible vials.

LC separation was performed on a Vanquish UHPLC System. Chromatography was carried out with an ACQUITY UPLC ® HSS T3 (150 × 2.1 mm, 1.8 μm). The column was maintained at 40°C. The flow rate and injection volume were set at 0.25 mL/min and 2 μL, respectively. For LC-ESI (+)-MS analysis, the mobile phases consisted of (B2) 0.1% formic acid in acetonitrile (v/v) and (A2) 0.1% formic acid in water (v/v). Separation was conducted under the following gradient: 0∼1 min, 2% B2; 1∼9 min, 2%∼50% B2; 9∼12 min, 50%∼98% B2; 12∼13.5 min, 98% B2; 13.5∼14 min, 98%∼2% B2; 14∼20 min, 2% B2. For LC-ESI (-)-MS analysis, the analysis was carried out with (B3) acetonitrile and (A3) ammonium formate (5 mM). Separation was conducted under the following gradient: 0∼1 min, 2% B3; 1∼9 min, 2%∼50% B3; 9∼12 min, 50%∼98% B3; 12∼13.5 min, 98% B3; 13.5∼14 min, 98%∼2% B3; 14∼17 min, 2% B3.

Mass spectrometric detection of metabolites was performed on Orbitrap Exploris 120 with an ESI ion source. Simultaneous MS1 and MS/MS (Full MS-ddMS2 mode, data-dependent MS/MS) acquisition was used. The parameters were as follows: sheath gas pressure, 30 arb; auxiliary gas flow, 10 arb; spray voltage, 3.50 kV and -2.50 kV for ESI (+) and ESI (-), respectively; capillary temperature, 325 °C; MS1 range, m/z 100-1000; MS1 resolving power, 60000 FWHM; number of data dependant scans per cycle, 4; MS/MS resolving power, 15000 FWHM; normalized collision energy, 30%; dynamic exclusion time, automatic.

Raw mass spectrometry downstream files were converted to mzXML file format by the MSConvert tool in the Proteowizard package (v3.0.8789). Peak detection, peak filtering, and peak alignment were performed using the R XCMS package to obtain a list of substances for quantification, with parameters set to bw=2, ppm=15, peakwidth=c(5, 30), mzwid=0.015, mzdiff=0.01, method="centWave ". Data correction was achieved by signal correction method based on QC samples to eliminate systematic errors, and substances with RSD > 30% in QC samples were filtered out for subsequent data analysis during the QC and QA processes.

Substance identification (library search) was performed using the public databases HMDB, massbank, LipidMaps, mzcloud, KEGG and the self-built standard library of PANOMIX metabolism, with parameters set to ppm < 30 ppm, to obtain the metabolite characterization results. The specific principle is to determine the molecular weight of the metabolite based on the mass-to-charge ratio (m/z) of the parent ion in the primary mass spectrometry, to predict the molecular formula by mass number deviation (ppm) and information such as addition ions, and then to match with the database to achieve the primary identification of the metabolite. Meanwhile, in the quantitative list, the metabolite detected in the secondary spectrum was matched with the fragment ion and other information of each metabolite in the database to achieve secondary identification of metabolites.

Sample data were subjected to principal component analysis (PCA), partial least squares discriminant analysis (PLS-DA), and orthogonal partial least squares discriminant analysis (OPLS-DA) dimensionality reduction analysis using the R package Ropls, respectively. R2X and R2Y indicate the explanatory rate of the proposed model to the X and Y matrices, respectively, and Q2 indicates the predictive power of the model; the closer their values are to 1, the better the fit of the model and the more accurately the samples in the training set can be classified into their original attribution. The strength and explanatory power of the influence of each metabolite content on the classification discrimination of samples were measured based on the Student’s t-test to calculate the nominal P-value, the OPLS-DA dimensionality reduction method to calculate the variable projection importance (VIP), and the fold change to calculate the component difference multiplier to aid in the screening of marker metabolites. Metabolite molecules were considered significantly different when the P-value < 0.05 and VIP value > 1.

#### • Targeted Metabolomic Sample Processing, Data Acquisition and Analysis

Cells were treated with creatine (10 mM) or distilled water (solvent control) for seven days. Cells were then washed twice with serine-free RPMI-1640 and seeded at 300,000 cells/mL with fresh complete media containing 30 mg/L of U^13^C-Serine 24 hours prior to harvesting. On the day of harvesting, cells were enumerated, washed with 1 mL PBS and stored as pellets at -80°C. A volume of 50 µL of spent media was also set aside for analysis. For detection of intracellular glycine, 7-day treated cells were seeded at 300,000 cells/mL in complete RPMI-1640 media and harvested after 8 hours, with the cells being pelleted and 50 µL of spent media being set aside for analysis.

Frozen cell pellets were extracted with dry ice-cold 80% methanol, sonicated for 3 minutes in ice-cold water, vortexed for 5 mins at 2000 rpm and 4°C, incubated on ice for 20 mins, centrifuged at 18,000g and 4°C for 20 minutes to collect the supernatant. The extract was dried down using the Genevac EZ-2.4 elite evaporator. On the day of analysis, the dried-down samples were reconstituted in an equal volume of 60:40 acetonitrile: water. Metabolites were separated using the Vanquish Horizon UHPLC system and iHILIC-(P) Classic (2.1x150 mm, 5 µm) columns. The high-resolution Orbitrap IQ-X Tribrid mass spectrometer with a H-ESI probe operating in switch polarity was used to detect and quantify the metabolite levels. The mobile phase A(MPA) was 20 mM ammonium bicarbonate at pH 9.6, adjusted by ammonium hydroxide addition and MPB was acetonitrile. The column temperature, injection volume, and the flow rate were 40°C, 2 µL, and 0.2mL/minute, respectively. The chromatographic gradient was 0 minutes: 85% B, 0.5 minutes: 85% B, 18 minutes: 20% B, 20 minutes: 20% B, 20.5 minutes: 85% B and 28 minutes: 85% B. MS parameters were as follows: Acquisition range of 70-1000 m/z at 60K resolution, spray voltage: 3600V for positive ionization and 2800 for negative ionization modes, sheath gas: 35, auxiliary gas: 5, sweep gas: 1, ion transfer tube temperature: 250°C, vaporizer temperature: 350°C, AGC target: 100%, and a maximum injection time of 118 ms. Poole QC samples were injected intermittently to determine the reproducibility of the detection and metabolite stability over time. Data acquisition was done using the ThermoFisher Xcalibur software and data analysis was performed using Skyline software^49^. Metabolite identification was done by matching the retention time and MS/MS fragmentation to the in-house database generated using the commercially available reference standards. Natural abundance correction was performed using the IsoCor^50^.

#### • Protein Electrophoresis and Immunoblotting

Cells were harvested for lysate preparation with protein levels normalized via the CellTiter-Glo® 2.0 Luminescence Assay. SDS-PAGE was subsequently carried out using 4–15% Mini-PROTEAN® TGX™ Precast Protein Gels and transferred onto a 0.45 µm PVDF membrane with a semi-dry transfer apparatus. The membrane was blocked with either 5% skim milk in TBST (1X TBS with 0.05% TWEEN® 20) for chemiluminescence imaging, or 1X BLOCKER solution for fluorescence imaging. Primary antibody incubation was performed at 4°C overnight with agitation. On the day of imaging, the membrane was washed thrice with TBST, incubated with secondary antibodies in appropriate diluents at room temperature for 30 minutes and washed thrice prior to imaging. Chemiluminescent imaging was performed with the ChemiDoc Imaging System. Protein detection utilized either the SuperSignal West Pico PLUS Chemiluminescent Substrate or the SuperSignal™ West Atto Ultimate Sensitivity Substrate depending on the level of expressed protein. Multiplex fluorescent imaging was performed with the iBright FL1500 Imaging System.

#### • Histological Staining, Data Acquisition and Analysis

Formalin-fixed paraffin-embedded PTLD tissue slices (3 μm) were dewaxed and serially stained for *EBER* RNA and GATM protein. Clinical-grade Epstein-Barr virus early RNA (EBER) ISH was performed by the Department of Pathology (National University Hospital) on a Bond III autostainer (Leica) using the Bond EBER probe together with BOND anti-fluorescein antibodies in accordance with ISH Protocol A in the manufacturer’s Instructions For Use (IFU). GATM immunohistochemical staining was performed according to the manufacturer’s protocol. Counterstaining was performed with hematoxylin. Images were captured using 40x air objective (Neo-Fluar numerical aperture 0.75) and stitched into whole sections using TissueFAXS Slide Scanner (TissueGnostics).

#### • Quantification and Statistical Analysis

Quantification and statistical analyses of all quantitative datasets were carried out with GraphPad Prism. FlowJo was used for analysis of flow cytometry data.

## Notes

### Competing Interest Statement

The authors have declared no competing interest.

## References

s1 . Opelz G, Döhler B. Lymphomas after solid organ transplantation: a collaborative transplant study report. Am J Transplant. Feb 2004;4(2):222–30. doi:10.1046/j.1600-6143.2003.00325.x

2. Wang LW, Shen H, Nobre L, et al. Epstein-Barr-Virus-Induced One-Carbon Metabolism Drives B Cell Transformation. Cell Metab. Sep 2019;30(3):539–555.e11. doi:10.1016/j.cmet.2019.06.003

3. Ma Y, Walsh MJ, Bernhardt K, et al. CRISPR/Cas9 Screens Reveal Epstein-Barr Virus-Transformed B Cell Host Dependency Factors. Cell Host Microbe. May 2017;21(5):580–591.e7. doi:10.1016/j.chom.2017.04.005

4. Di Pietro E, Sirois J, Tremblay ML, MacKenzie RE. Mitochondrial NAD-dependent methylenetetrahydrofolate dehydrogenase-methenyltetrahydrofolate cyclohydrolase is essential for embryonic development. Mol Cell Biol. Jun 2002;22(12):4158–66.

5. Ron-Harel N, Santos D, Ghergurovich JM, et al. Mitochondrial Biogenesis and Proteome Remodeling Promote One-Carbon Metabolism for T Cell Activation. Cell Metab. Jul 2016;24(1):104–17. doi:10.1016/j.cmet.2016.06.007

6. Ma EH, Bantug G, Griss T, et al. Serine Is an Essential Metabolite for Effector T Cell Expansion. Cell Metab. Feb 7 2017;25(2):345–357. doi:10.1016/j.cmet.2016.12.011

7. Nilsson R, Jain M, Madhusudhan N, et al. Metabolic enzyme expression highlights a key role for MTHFD2 and the mitochondrial folate pathway in cancer. Nat Commun. 2014;5:3128. doi:10.1038/ncomms4128

8. Gustafsson Sheppard N, Jarl L, Mahadessian D, et al. The folate-coupled enzyme MTHFD2 is a nuclear protein and promotes cell proliferation. Sci Rep. 2015;5:15029. doi:10.1038/srep15029

9. Koufaris C, Nilsson R. Protein interaction and functional data indicate MTHFD2 involvement in RNA processing and translation. Cancer Metab. 2018;6:12. doi:10.1186/s40170-018-0185-4

10. Ducker GS, Chen L, Morscher RJ, et al. Reversal of Cytosolic One-Carbon Flux Compensates for Loss of the Mitochondrial Folate Pathway. Cell Metab. Jun 2016;23(6):1140–53. doi:10.1016/j.cmet.2016.04.016

11. Baker SA, Gajera CR, Wawro AM, Corces MR, Montine TJ. GATM and GAMT synthesize creatine locally throughout the mammalian body and within oligodendrocytes of the brain. Brain Res. Nov 01 2021;1770:147627. doi:10.1016/j.brainres.2021.147627

12. Jiang S, Wang LW, Walsh MJ, et al. CRISPR/Cas9-Mediated Genome Editing in Epstein-Barr Virus-Transformed Lymphoblastoid B-Cell Lines. Curr Protoc Mol Biol. Jan 2018;121:31.12.1-31.12.23. doi:10.1002/cpmb.51

13. Minton DR, Nam M, McLaughlin DJ, et al. Serine Catabolism by SHMT2 Is Required for Proper Mitochondrial Translation Initiation and Maintenance of Formylmethionyl-tRNAs. Mol Cell. Feb 15 2018;69(4):610–621.e5. doi:10.1016/j.molcel.2018.01.024

14. Gewurz B, Guo R, Lim M, et al. Multi-omic Analysis of Human B-cell Activation Reveals a Key Lysosomal BCAT1 Role in mTOR Hyperactivation by B-cell receptor and TLR9. Res Sq. May 30 2024;doi:10.21203/rs.3.rs-4413958/v1

15. Arvey A, Tempera I, Tsai K, et al. An atlas of the Epstein-Barr virus transcriptome and epigenome reveals host-virus regulatory interactions. Cell Host Microbe. Aug 2012;12(2):233–45. doi:10.1016/j.chom.2012.06.008

16. Smith GK, Mueller WT, Wasserman GF, Taylor WD, Benkovic SJ. Characterization of the enzyme complex involving the folate-requiring enzymes of de novo purine biosynthesis. Biochemistry. Sep 02 1980;19(18):4313–21. doi:10.1021/bi00559a026

17. WALKER JB. Repression of arginine-glycine transamidinase activity by dietary creatine. Biochim Biophys Acta. Dec 1959;36:574–5. doi:10.1016/0006-3002(59)90217-3

18. Derave W, Marescau B, Vanden Eede E, Eijnde BO, De Deyn PP, Hespel P. Plasma guanidino compounds are altered by oral creatine supplementation in healthy humans. J Appl Physiol (1985). Sep 2004;97(3):852-7. doi:10.1152/japplphysiol.00206.2004

19. Jiang S, Zhou H, Liang J, et al. The Epstein-Barr Virus Regulome in Lymphoblastoid Cells. Cell Host Microbe. Oct 2017;22(4):561–573.e4. doi:10.1016/j.chom.2017.09.001

20. Gires O, Zimber-Strobl U, Gonnella R, et al. Latent membrane protein 1 of Epstein-Barr virus mimics a constitutively active receptor molecule. EMBO J. Oct 1997;16(20):6131–40. doi:10.1093/emboj/16.20.6131

21. Uchida J, Yasui T, Takaoka-Shichijo Y, et al. Mimicry of CD40 signals by Epstein-Barr virus LMP1 in B lymphocyte responses. Science. Oct 1999;286(5438):300-3.

22. Oton AB, Wang H, Leleu X, et al. Clinical and pathological prognostic markers for survival in adult patients with post-transplant lymphoproliferative disorders in solid transplant. Leuk Lymphoma. Sep 2008;49(9):1738–44. doi:10.1080/10428190802239162

23. Uhlen M, Zhang C, Lee S, et al. A pathology atlas of the human cancer transcriptome. Science. Aug 18 2017;357(6352)doi:10.1126/science.aan2507

24. Expression of GATM in renal cancer - The Human Protein Atlas. The Human Protein Atlas. https://www.proteinatlas.org/ENSG00000171766-GATM/cancer/renal+cancer

25. Expression of GAMT in renal cancer - The Human Protein Atlas. The Human Protein Atlas. https://www.proteinatlas.org/ENSG00000130005-GAMT/cancer/renal+cancer

26. Yang A, Guo L, Zhang Y, et al. MFN2-mediated mitochondrial fusion facilitates acute hypobaric hypoxia-induced cardiac dysfunction by increasing glucose catabolism and ROS production. Biochim Biophys Acta Gen Subj. Sep 2023;1867(9):130413. doi:10.1016/j.bbagen.2023.130413

27. Faulkner MC, Miller DW, Hatch GM, inventors; Vireo Systems, Inc., Madison, TN, USA, assignee. Compositions and methods for neuroprotection and treatment of neurodegeneration. 2021.

28. Borsook H, Dubnoff JW. THE FORMATION OF CREATINE FROM GLYCOCYAMINE IN THE LIVER. Journal of Biological Chemistry. 1940/02/01/ 1940;132(2):559-574. 10.1016/S0021-9258(19)56203-2

29. Borsook H, Dubnoff JW. THE FORMATION OF GLYCOCYAMINE IN ANIMAL TISSUES. Journal of Biological Chemistry. 1941/03/01/ 1941;138(1):389-403. 10.1016/S0021-9258(18)51446-0

30. Horner WH. Transamidination in the Nephrectomized Rat. Journal of Biological Chemistry. 1959/09/01/ 1959;234(9):2386–2387. 10.1016/S0021-9258(18)69820-5

31. Jain M, Nilsson R, Sharma S, et al. Metabolite profiling identifies a key role for glycine in rapid cancer cell proliferation. Science. May 25 2012;336(6084):1040-4. doi:10.1126/science.1218595

32. Meiser J, Tumanov S, Maddocks O, et al. Serine one-carbon catabolism with formate overflow. Sci Adv. Oct 2016;2(10):e1601273. doi:10.1126/sciadv.1601273

33. Xu Y, Ren J, Wang W, Zeng AP. Improvement of glycine biosynthesis from one-carbon compounds and ammonia catalyzed by the glycine cleavage system in vitro. Eng Life Sci. Jan 2022;22(1):40–53. doi:10.1002/elsc.202100047

34. Leung KY, De Castro SCP, Galea GL, Copp AJ, Greene NDE. Glycine Cleavage System H Protein Is Essential for Embryonic Viability, Implying Additional Function Beyond the Glycine Cleavage System. Front Genet. 2021;12:625120. doi:10.3389/fgene.2021.625120

35. Lunetti P, Damiano F, De Benedetto G, et al. Characterization of Human and Yeast Mitochondrial Glycine Carriers with Implications for Heme Biosynthesis and Anemia. J Biol Chem. Sep 16 2016;291(38):19746–59. doi:10.1074/jbc.M116.736876

36. Fritsche E, Humm A, Huber R. Substrate binding and catalysis by L-arginine:glycine amidinotransferase--a mutagenesis and crystallographic study. Eur J Biochem. Jul 15 1997;247(2):483–90. doi:10.1111/j.1432-1033.1997.00483.x

37. Gross MD, Eggen MA, Simon AM, Van Pilsum JF. The purification and characterization of human kidney L-arginine:glycine amidinotransferase. Arch Biochem Biophys. Dec 1986;251(2):747–55. doi:10.1016/0003-9861(86)90385-1

38. Hariharan S, McBride MA, Cherikh WS, Tolleris CB, Bresnahan BA, Johnson CP. Post- transplant renal function in the first year predicts long-term kidney transplant survival. Kidney Int. Jul 2002;62(1):311–8. doi:10.1046/j.1523-1755.2002.00424.x

39. Doench JG, Fusi N, Sullender M, et al. Optimized sgRNA design to maximize activity and minimize off-target effects of CRISPR-Cas9. Nat Biotechnol. Feb 2016;34(2):184–91. doi:10.1038/nbt.3437

40. Wang LW, Wang Z, Ersing I, et al. Epstein-Barr virus subverts mevalonate and fatty acid pathways to promote infected B-cell proliferation and survival. PLoS Pathog. 09 2019;15(9):e1008030. doi:10.1371/journal.ppat.1008030

41. Picelli S, Faridani OR, Björklund AK, Winberg G, Sagasser S, Sandberg R. Full-length RNA-seq from single cells using Smart-seq2. Nat Protoc. Jan 2014;9(1):171–81. doi:10.1038/nprot.2014.006

42. Andrews S. FastQC: A Quality Control tool for High Throughput Sequence Data. Babraham Institute. https://www.bioinformatics.babraham.ac.uk/projects/fastqc/

43. Ewels P, Magnusson M, Lundin S, Käller M. MultiQC: summarize analysis results for multiple tools and samples in a single report. Bioinformatics. Oct 01 2016;32(19):3047–8. doi:10.1093/bioinformatics/btw354

44. Dobin A, Davis CA, Schlesinger F, et al. STAR: ultrafast universal RNA-seq aligner. Bioinformatics. Jan 01 2013;29(1):15–21. doi:10.1093/bioinformatics/bts635

45. Liao Y, Smyth GK, Shi W. featureCounts: an efficient general purpose program for assigning sequence reads to genomic features. Bioinformatics. Apr 01 2014;30(7):923–30. doi:10.1093/bioinformatics/btt656

46. Harrow J, Denoeud F, Frankish A, et al. GENCODE: producing a reference annotation for ENCODE. Genome Biol. 2006;7 Suppl 1(Suppl 1):S4.1-9. doi:10.1186/gb-2006-7-s1-s4

47. Robinson MD, McCarthy DJ, Smyth GK. edgeR: a Bioconductor package for differential expression analysis of digital gene expression data. Bioinformatics. Jan 01 2010;26(1):139–40. doi:10.1093/bioinformatics/btp616

48. Weekes MP, Tomasec P, Huttlin EL, et al. Quantitative temporal viromics: an approach to investigate host-pathogen interaction. Cell. Jun 2014;157(6):1460–72. doi:10.1016/j.cell.2014.04.028

49. Pino LK, Searle BC, Bollinger JG, Nunn B, MacLean B, MacCoss MJ. The Skyline ecosystem: Informatics for quantitative mass spectrometry proteomics. Mass Spectrom Rev. May 2020;39(3):229–244. doi:10.1002/mas.21540

50. Millard P, Delépine B, Guionnet M, Heuillet M, Bellvert F, Létisse F. IsoCor: isotope correction for high-resolution MS labeling experiments. Bioinformatics. Nov 01 2019;35(21):4484–4487. doi:10.1093/bioinformatics/btz209

