## Supplemental Figures and Tables for "Creatine synthesis is a tumor suppressor pathway hypostatic to one-carbon metabolism"

**Figure S1, related to Figure 1**

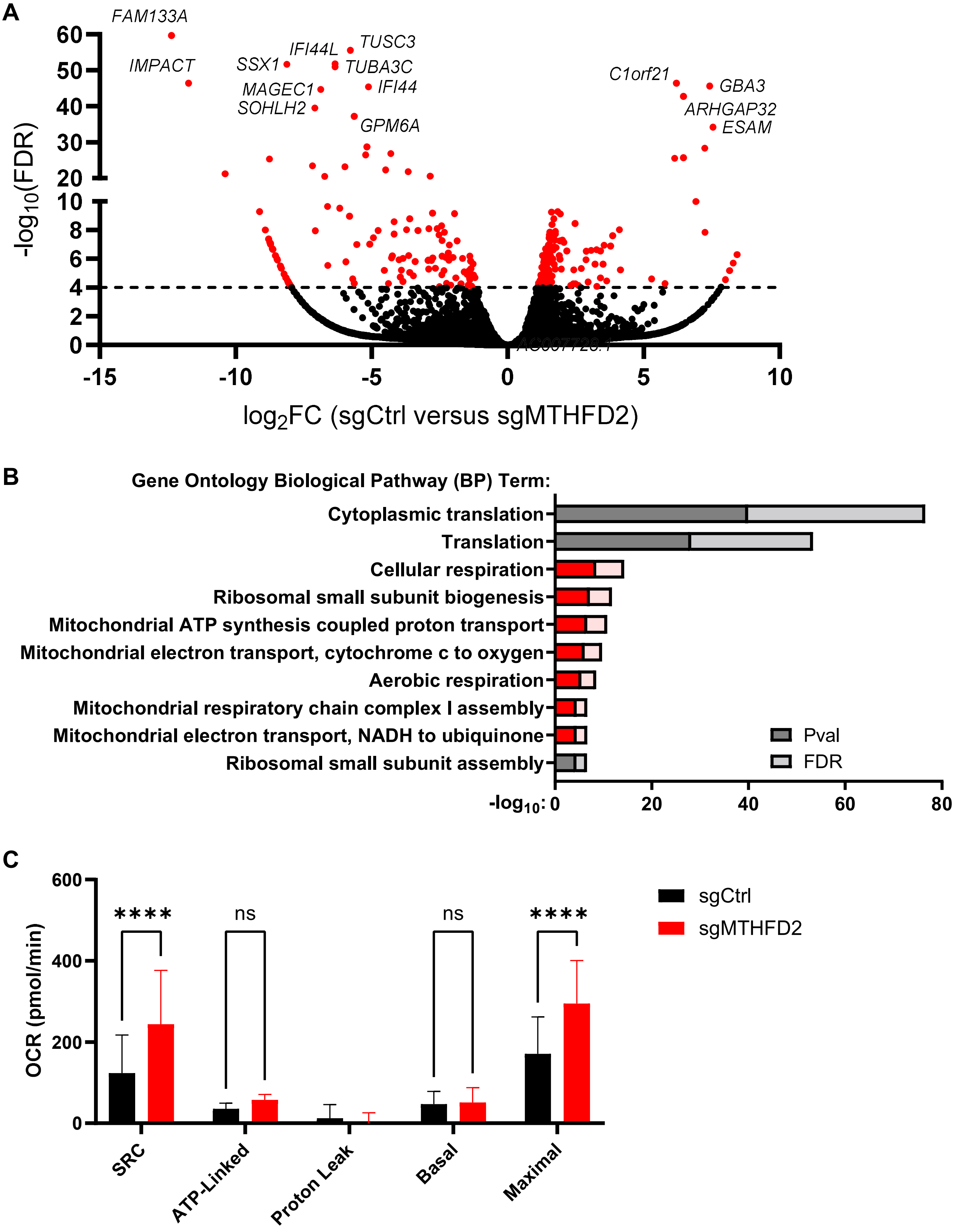

**Figure S1 Loss of MTHFD2 remodels the transcriptome and enhances oxidative capacity.**

(A) Volcano plot illustrating the differentially regulated genes in GM12878 sgCtrl and sgMTHFD2 cells. The Y-intercept indicates the false discovery rate (FDR) cut-off of 0.0001. Red dots represent genes that are significantly different between sgCtrl and sgMTHFD2. Labeled dots indicate genes that are significantly regulated i.e. $log_2FC > 5$ or $log_2FC < -5$.

(B) Gene Ontology analysis of significantly regulated genes to determine biological pathway enrichment. Dark grey bars represent nominal P-values while light grey bars represent FDR values. Bars corresponding to BP terms related to oxidative phosphorylation (OXPHOS) are indicated in red (representing nominal P-values; counterpart to dark grey bars) and pink (representing FDR values; counterpart to light grey bars).

(C) Detailed breakdown of OXPHOS parameters measured from the Seahorse XF Cell Mito Stress Test. Data represent N = 3 biological replicates with errors bars indicative of SD. ns, not significant; ****, $p < 0.0001$ by Student's t-test.

### Figure S2, related to Figure 2

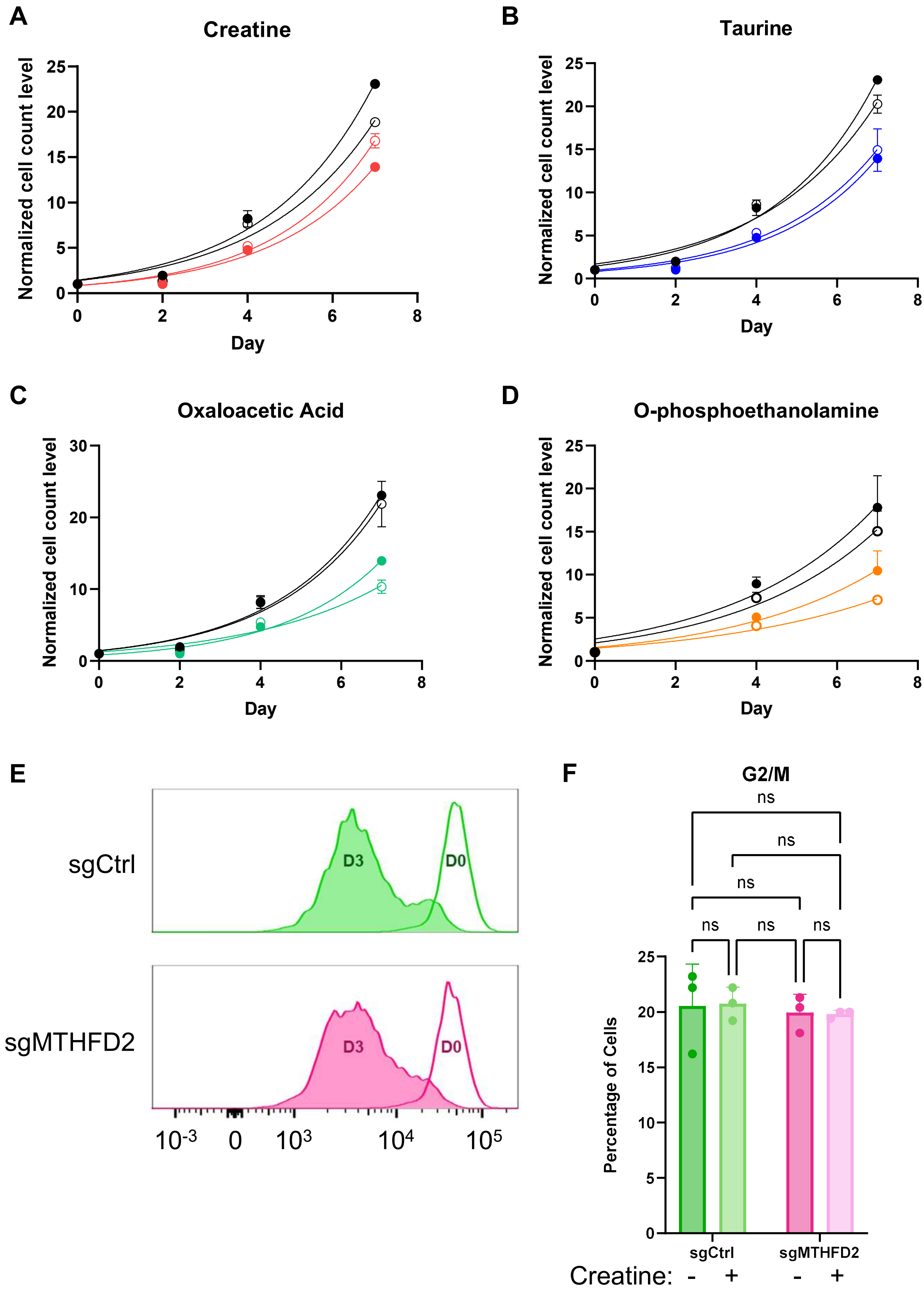

**Figure S2 Viability defects in MTHFD2 ablation are not due to proliferative or mitotic arrest and can be mitigated with creatine supplementation.**

Relative cell numbers of GM12878 sgCtrl and sgMTHFD2 cells supplemented with various metabolites: (A) creatine, (B) taurine, (C) oxaloacetic acid, (D) O-phosphoethanolamine. Circles are denoted as follows: black filled = sgCtrl + water; black empty = sgCtrl + supplement; colored filled = sgMTHFD2 + water; colored empty = sgMTHFD2 + supplement. Data represent N = 3 biological replicates with errors bars indicative of SD.

(E) Representative histograms (N = 3) of GM12878 sgCtrl and sgMTHFD2 stained with CellTrace Violet dye to track proliferation by dye dilution assay. D0, day 0 i.e., day of cell staining and seeding; D3, day 3 post-seeding. Histograms were transformed by normalization to mode.

(F) Frequency plots of the G2/M subpopulation from the cell cycle analysis shown in Figure 2C. -, absence of supplement; +, presence of supplement. Data represent N = 3 biological replicates with errors bars indicative of SD. ns, not significant by ordinary two-way ANOVA with uncorrected Fisher's LSD.

**Figure S3, related to Figure 3**

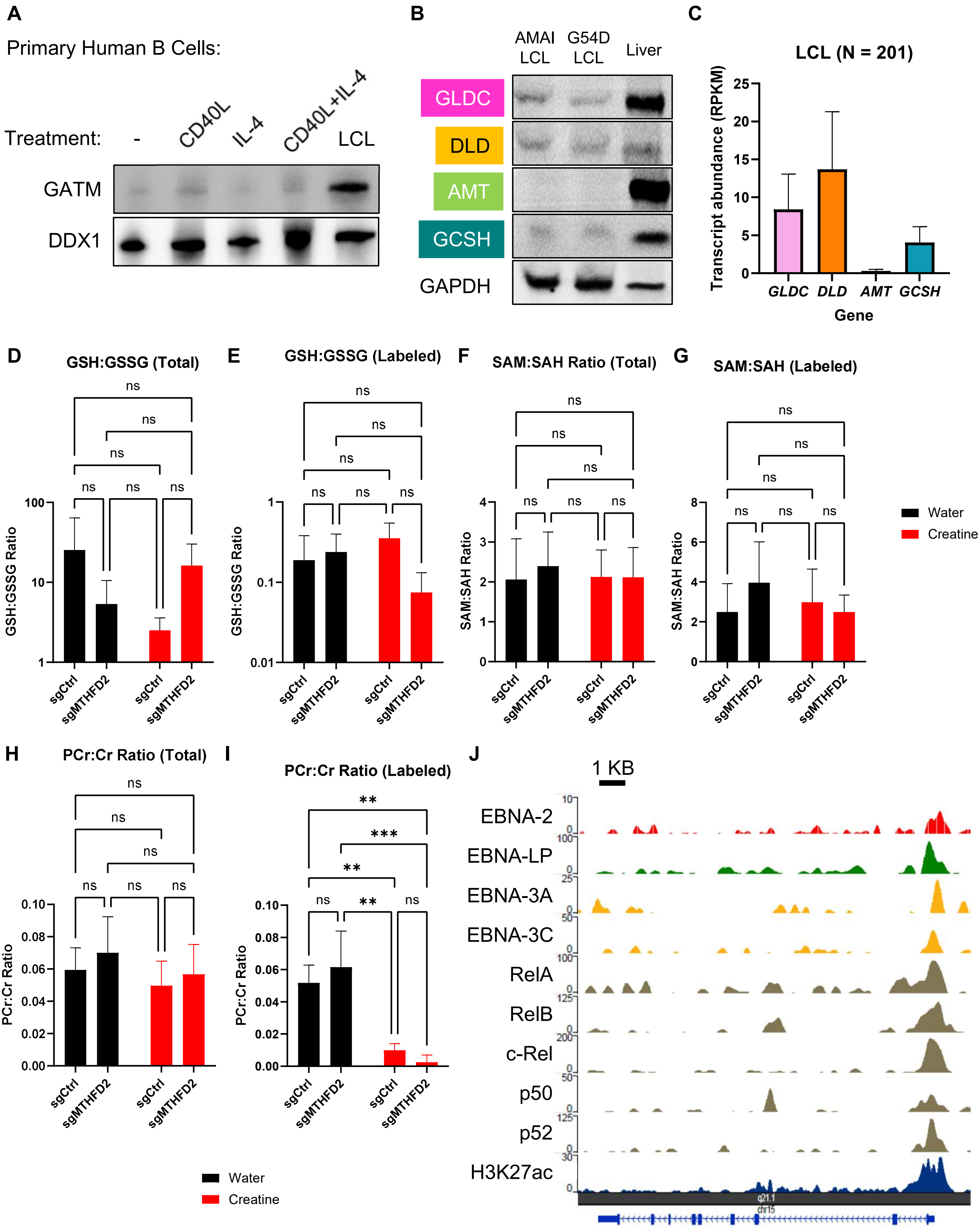

**Figure S3 Creatine synthesis is uniquely induced by EBV in LCL, and LCL has a short-circuited glycine cleavage system.**

(A) Representative immunoblots (N = 3) of samples from primary human B cells stimulated with either CD40 ligand (CD40L), interleukin-4 (IL-4) or both, as well as G54D LCL. GATM was probed for as the protein of interest. DDX1 was used as the housekeeping gene control.

(B) Representative immunoblots (N = 3) of LCL and normal human liver samples. Enzymes of the glycine cleavage system (GCS), namely GLDC, DLD, AMT and GCSH, were probed for as the proteins of interest. GAPDH was used as the housekeeping gene control.

(C) Plot of transcript abundances for various GCS genes across a set of public LCL transcriptomes. Data represent N = 201 biologically distinct lines with error bars indicative of SD.

Measurements of metabolite pairs in GM12878 sgCtrl and sgMTHFD2 cells treated with either water (control; denoted by black bars) or 10 mM creatine (denoted by red bars), in either totality or the $^{13}$C-labeled fractions. Metabolite pairs analyzed were (D) Total GSH:GSSG, (E) Labeled GSH:GSSG, (F) Total SAM:SAH, (G) Labeled SAM:SAH, (H) Total phospho-creatine (PCr):creatine (Cr), (I) Labeled PCr:Cr. Data represent N = 3 biological replicates with errors bars indicative of SD. ns, not significant; **, $p < 0.01$; ***, $p < 0.001$ by ordinary two-way ANOVA with uncorrected Fisher's LSD.

(J) Plot of public chromatin immunoprecipitation (ChIP)-seq tracks at the *GATM* locus showing the colocalization of EBV and NF-κB factors at the promoter.

**Figure S4, related to Figure 4**

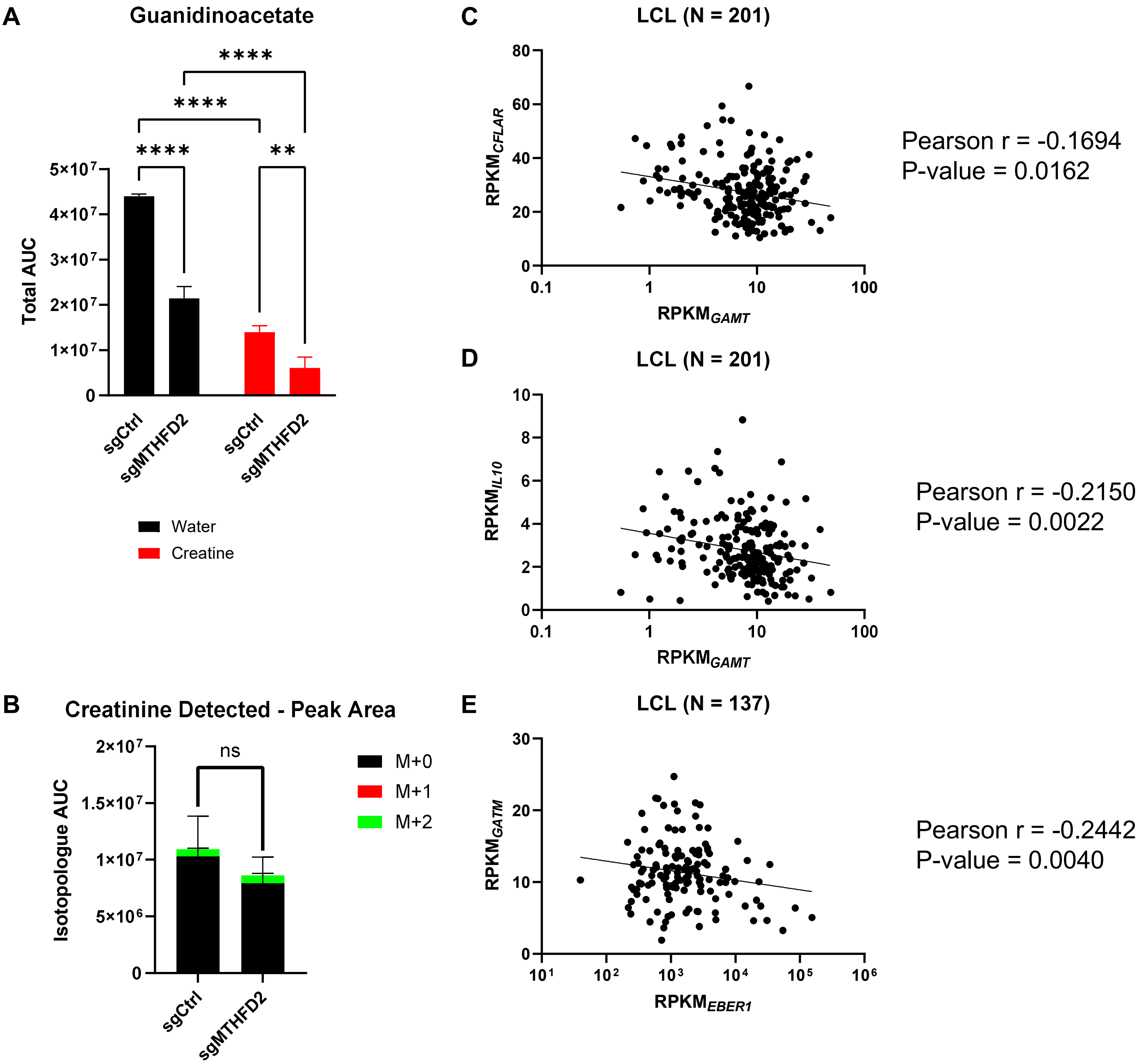

**Figure S4 Regulation of cellular creatine and creatine's correlation against PTLD-driving molecules.**

(A) Levels of intracellular guanidinoacetate in GM12878 sgCtrl and sgMTHFD2 cells treated with either water (control; denoted by black bars) or 10 mM creatine (denoted by red bars). AUC, area under curve. Data represent N = 3 biological replicates with errors bars indicative of SD. **, $p < 0.01$; ****, $p < 0.0001$ by ordinary two-way ANOVA with uncorrected Fisher's LSD.

(B) Isotopologue abundances for creatinine in GM12878 sgCtrl and sgMTHFD2 cells. Data represent N = 3 biological replicates with errors bars indicative of SD. ns, not significant by Student's t-test.

Correlation analysis of selected genes in public LCL transcriptomes, namely (C) *CFLAR* against *GAMT* (N = 201), (D) *IL10* against *GAMT* (N = 201), and (E) *GATM* against *EBER1* (N = 137). For statistical testing, Pearson correlation coefficients were computed with the exact P-values indicated.

**Table S1 PTLD patient details.**

| Patient ID | Age (years) | Sex | PTLD Type | Organ involved | Previous Solid Organ Transplant, Vintage of Transplant (years, if known) |
|---|---|---|---|---|---|
| P1 | 5 | Female | Polymorphic | Tonsil | Liver, 2 |
| P2 | 4 | Male | Polymorphic | Stomach, intestine | Liver, 1 |
| P3 | 2 | Female | Polymorphic | Stomach, intestine | Liver |
| P4 | 25 | Male | Monomorphic | Tonsil | Kidney, <1 |
| P5 | 11 | Female | Monomorphic | Gum | Liver |
| P6 | 2 | Male | Monomorphic | Stomach, intestine | Liver |
| C1 | 2 | Male | Control | Tonsil | NA |
| C2 | 4 | Female | Control | Tonsil | NA |
| C3 | 4 | Male | Control | Tonsil | NA |

Relevant patient details are shown here. P2 and P6 are illustrated in Figure 4D as the polymorphic and monomorphic PTLD samples, respectively. Of the available cohort, these two patients had similar demographic and clinical characteristics, and were therefore the best pair for direct comparison.
